# Karyopherin mimicry explains how the HIV capsid penetrates nuclear pores

**DOI:** 10.1101/2023.03.23.534032

**Authors:** C.F. Dickson, S. Hertel, J. Ruan, N. Ariotti, A. Tuckwell, N. Li, S.C. Al-Izzi, E. Sierecki, Y. Gambin, R.G. Morris, G.J. Towers, T. Böcking, D.A. Jacques

**Affiliations:** UNSW Sydney; The University of New South Wales; The University of Queensland; University of New South Wales; EMBL Australia, The University of New South Wales; University College London

## Abstract

HIV can infect non-dividing cells because the viral capsid can overcome the selective barrier of the nuclear pore complex and deliver the genome directly into the nucleus. Remarkably, the intact HIV capsid is over one thousand times greater than the size-limit prescribed by the nuclear pore’s diffusion barrier. This barrier is a phase-separated condensate in the central channel of the nuclear pore and is comprised of intrinsically-disordered nucleoporin domains enriched in phenylalanine-glycine (FG) dipeptides. Through multivalent FG-interactions, cellular karyopherins and their bound cargoes solubilise in this phase to drive nucleocytoplasmic transport. By performing an *in vitro* dissection of the nuclear pore complex, we show that a pocket on the surface of the HIV capsid similarly interacts with FG-motifs from multiple nucleoporins and that this interaction licenses capsids to penetrate nucleoporin condensates. This karyopherin mimicry model resolves a key conceptual challenge for the role of the HIV capsid in nuclear entry, and explains how an exogenous entity much larger than any known cellular cargo can non-destructively breach the nuclear envelope.

## Introduction

All retroviruses establish infection by integrating their DNA into host chromatin in the nucleus. For ten of the eleven retrovirus genera, integration occurs during mitosis when the nuclear envelope disintegrates and the chromosomal DNA is exposed. However, lentiviruses, which include HIVs, have evolved to infect non-dividing cells. To do so, these viruses must navigate the gatekeeper of nuclear entry, the Nuclear Pore Complex (NPC).

The human NPC is a 110 MDa complex, comprised of 30 different nucleoporin proteins (Nups), which occur in stoichiometries ranging from 8-48 copies per NPC^1–4^. Movement through the NPC central transport channel is restricted by size. The vast majority of proteins larger than ∼40 kDa must recruit karyopherins to facilitate their nuclear transport^5, 6^. The selective channel gating nuclear entry is filled with a distinct phase-separated ‘gel’ formed from intrinsically disordered Nup domains that are enriched in Phenylalanine-Glycine (FG)-motifs. These FG-repeat domains are found in approximately one third of the Nups (FG-Nups), and contribute in excess of 5800 individual FG motifs to a single NPC.

HIV’s ability to infect non-dividing cells maps to the capsid protein (CA)^7, 8^. CA self-assembles into a metastable lattice to form a closed ‘shell’ of ∼40 MDa which contains the genomic RNA and viral enzymes. Although originally thought to disassemble shortly after entry into the host cell, it is now broadly accepted that the integrity of the CA lattice is vital for cellular trafficking, reverse transcription, and for protecting the viral genome from cytoplasmic nucleic acid sensors and nucleases^9–12^. Furthermore, recent imaging studies have captured intact capsids as they transit the NPC^13^ and have demonstrated intact capsids within the nucleus^14–18^.

Critically, it has been unclear how the HIV capsid can overcome the NPC’s selectivity barrier when it is over one thousand times greater in size than the passive diffusion limit. Transporters in the karyopherin family can carry large cargoes through the NPC by specifically interacting with the FG motifs of the gel-like diffusion barrier. The efficiency of this cargo transport^19–21^ relies on the presence of multiple FG-binding sites on the karyopherin. The interactions are highly specific, but weak and with rapid exchange kinetics, enabling the karyopherin and its bound cargo to partition into the selectivity barrier and rapidly diffuse through the NPC^22–25^. Co-crystal structures have shown that the HIV capsid exterior also possesses an FG-binding pocket, which recruits the cellular cofactors Sec24C^26^, Nup153^27^, and cleavage polyadenylation specific factor 6 (CPSF6)^28, 29^. While these proteins are structurally distinct and found in different cellular compartments, each of their interactions with the HIV capsid depends on an FG-motif which buries itself into a pocket in the CA N-terminal domain.

### HIV capsids bind to FG-repeat nucleoporins

Given that each CA molecule carries an FG-binding site, complete capsids carry over 1200 such sites and therefore have a high capacity to interact with proteins carrying multiple FG-motifs. We hypothesised that many CA:FG-Nup interactions occur in addition to those previously described and that even weak interactions at this site have the potential to be significant due to avidity affects. We recently developed an *in vitro* technique based on fluorescence fluctuation spectroscopy (FFS) to screen for interactions between the HIV capsid and cellular proteins^30^. Briefly, putative binders are expressed as GFP-fusions in a cell-free system. Upon addition of fluorescent capsid like particles (CLPs; CA_A204C_ cross-linked assemblies labelled with Alexa Fluor 568) binding is determined by two colour coincidence detection as individual CLPs pass through a confocal volume generating fluorescence fluctuations. Observations are made directly in the cell-free extract, which we have found to be a powerful approach to screening difficult-to-purify putative capsid binders such as the highly disordered FG-Nups. Furthermore, the use of CLPs as the binding platform has three critical advantages: (i) CLPs display a mix of cone, tube, and sphere morphologies bearing both CA hexamers and pentamers, and therefore they present all possible CA interfaces; (ii) the presence of a CA lattice gives high sensitivity to the measurement by physically concentrating binders on the CLP, thereby enabling detection of interactions well below the *K_D_*; and (iii) the CLP lattice can engage binders that rely on multivalent CA contacts to make high avidity capsid interactions.

To ‘dissect’ the NPC we individually expressed GFP-fusions of the 10 FG-repeat NPC components (Nup50, Nup54, Nup58, Nup62, Nup42, Nup98, Nup153, Nup214, Nup358 and Pom121; each with 5-42 FG-motifs) along with Nup35, Nup88, and Nup133, which do not have FG-repeat domains (for details see Table S1). We also included known CA-binders CPSF6 and CypA. For Nups that did not express as full-length proteins, we expressed only the FG-domain (Nup98_1-499_, Nup214_1210-2090_). Nups either remained as monomeric protein (exemplified by POM121, Fig 1a, flat blue trace), or spontaneously oligomerised (exemplified by Nup98, Fig 1b, fluctuating blue trace). This latter behaviour was not unexpected, given that FG-repeat proteins are known to phase-separate^31, 32^. Binding was identified by the recording of ‘mirror plots’ upon addition of CLP (Fig 1 a,b and Fig S1, orange traces, represented as negative values for clarity), showing that capsid and Nup were collocated. The degree of coincidence is reported as the ‘Nup:CA intensity ratio’^30^. The positive controls (CPSF6 and CypA) were recruited to the CA lattice as expected (Fig S1), while the non-FG-repeat Nups (Nup35, Nup88, and Nup133) showed no detectable binding (Fig 1c,d and Fig S1). Of the monomeric Nups, those with the highest number of FG motifs in unstructured FG domains, Nup58 (14 FGs), Pom121 (24 FGs) and Nup214 (42 FGs), showed clear CLP binding (Fig 1c and Fig S1). A similar trend was observed for the oligomeric Nups, with Nup42 (12 FGs), Nup153 (29 FGs) and Nup98 (40 FGs) all binding (Fig 1d and Fig S1). When accounting for stoichiometry, the six Nups that displayed CLP binding contribute 78% of the total FG repeat motifs in the NPC (Fig 1e). Furthermore, these Nups are found distributed throughout the NPC including in the cytoplasmic filaments (Nups 42, 98, and 214), the central transport channel (Nup58, Nup98 and POM121), and the nuclear basket (Nup153) (Fig 1 f). The clear coincidence between CLPs and Nup98 was particularly striking as this nucleoporin is present at 48 copies per NPC making it the most significant contributor of FGs to the diffusion barrier.

**Fig. 1.**
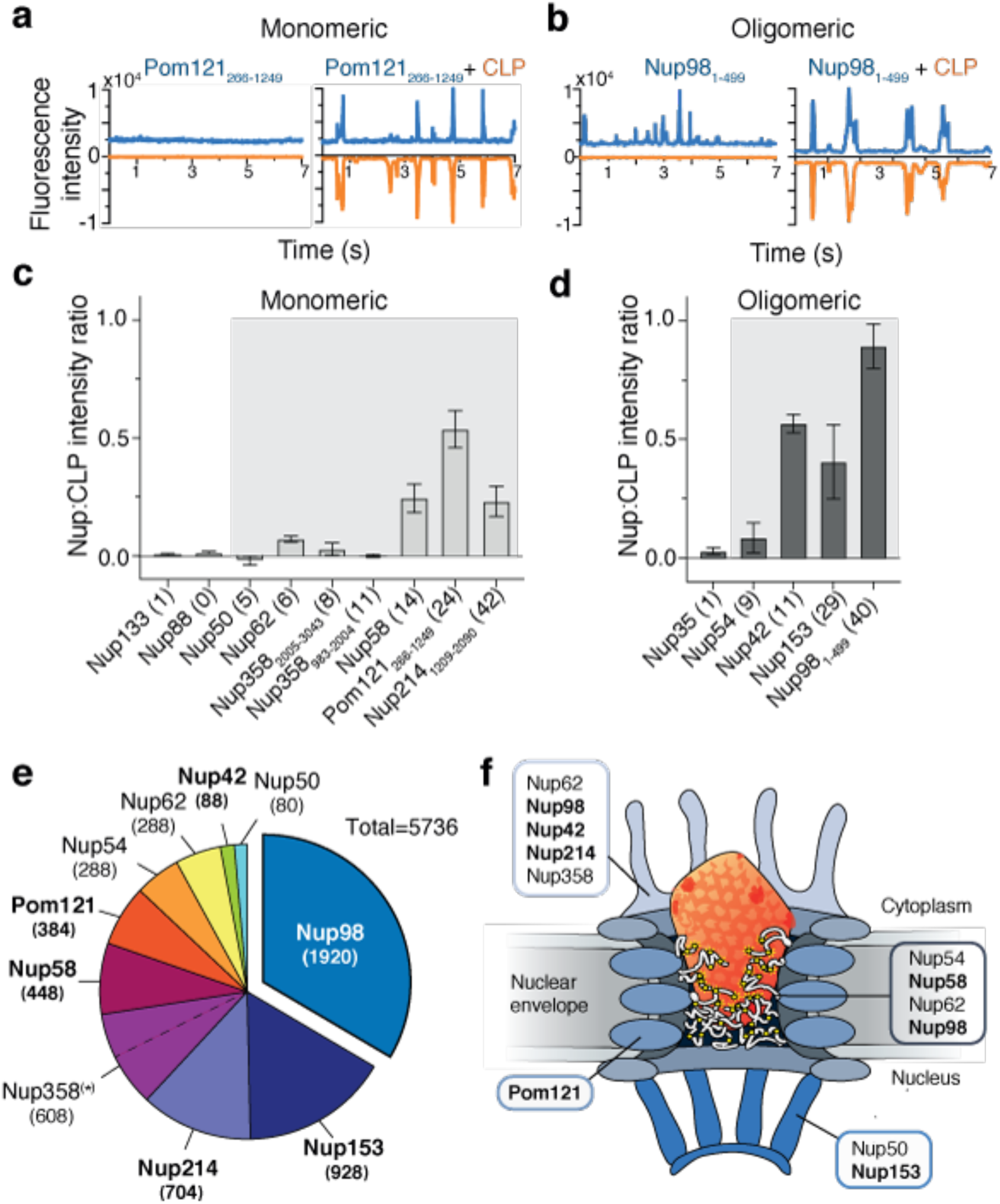
FG-Nups bind to the HIV-1 capsid. **a,b**, Fluorescence fluctuation spectroscopy traces of GFP-Pom121 (a) or GFP-Nup98 (b), alone or mixed with AF568 CLPs (blue, Nup channel; orange, CLP channel). **c,d**, Nup:CLP fluorescence intensity ratios calculated for monomeric (c) or oligomeric (d) Nups (grey background denotes canonical FG-repeat Nups with the number of FG-motifs per Nup in parentheses). Error bars are standard deviation. **e**, Relative FG-motif contributions to the diffusion barrier based on published Nup stoichiometries (see also Table S1). **f**, HIV capsid interacts with Nups distributed throughout the NPC. Nups identified to bind to CLPs are highlighted in bold.

### FG-repeat domains are specifically recruited to the FG-binding pocket on the HIV capsid

FG-containing peptides are known to have different capsid binding modes. Previous crystals structures have shown that the short peptides Nup153_1407-1423_ and CPSF6_313-327_ bury their respective FG motifs in the CA N-terminal domain. While these FG-motifs are conformationally identical, the peripheral interactions differ. Nup153_1407-1423_ adopts a linear conformation that bridges two CA monomers, while CPSF6_313-327_ forms a more compact structure which packs predominantly against a single CA protomer (Fig 2a). Importantly, in both cases CA residue N57 forms two critical hydrogen bonds with the mainchain NH and O of the buried phenylalanine. Mutation of CA N57 therefore disrupts binding of both cofactors. However, mutation of residues N74 or A77 only disrupts CPSF6 binding leaving Nup153 unaffected^28^. CLP mutants (N57D, N74D, or A77V) reproduced these binding specificities in our FFS assay (Fig 2 b,c and Fig S2-1), while binding of a CypA control was unaffected by these mutations, demonstrating these CLPs to be otherwise assembly- and binding-competent (Fig S2-1).

**Fig. 2.**
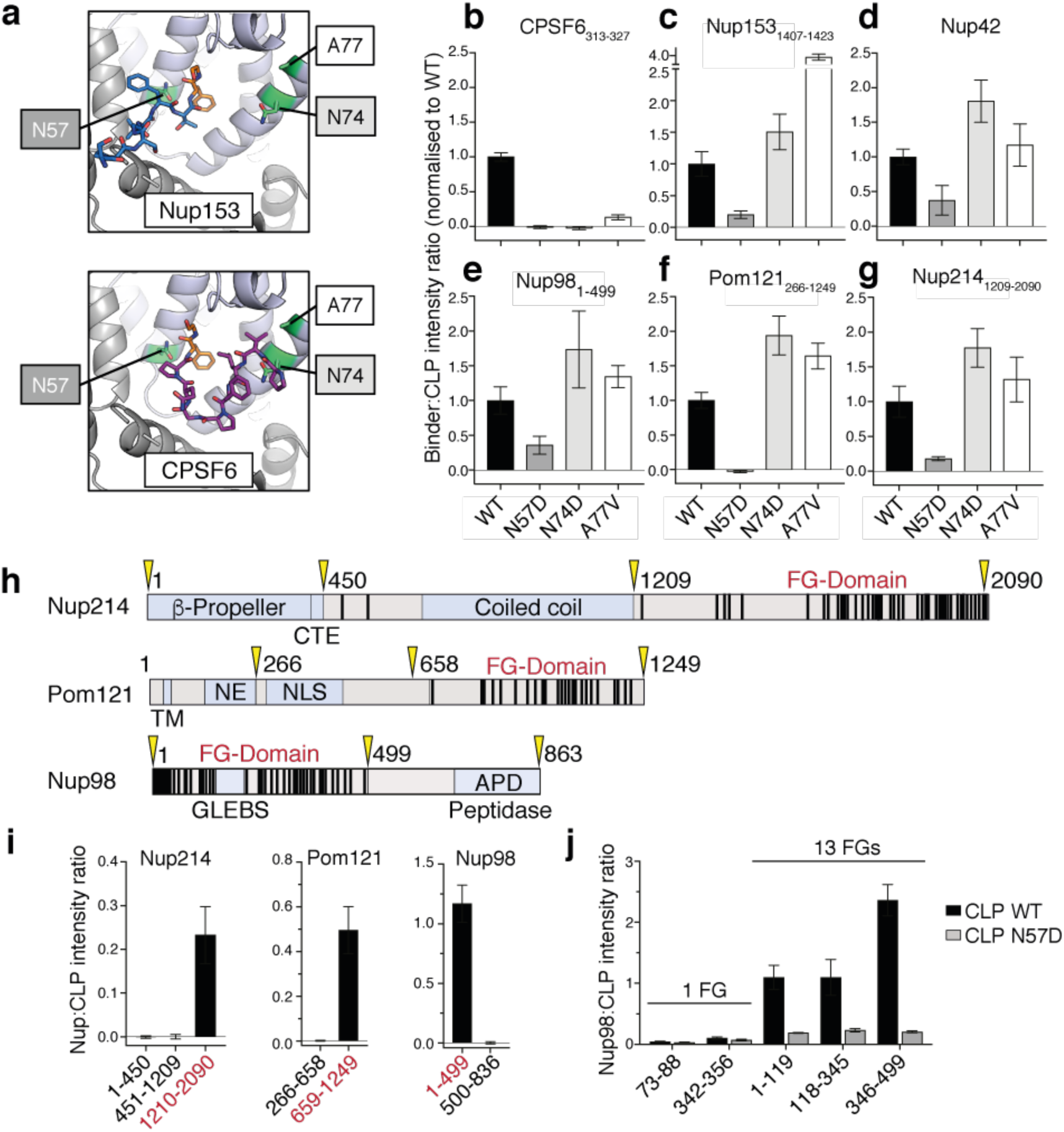
HIV-1 CA specifically recruits FG-repeat domains. **a**, Hydrophobic pocket of CA (cartoon) with Nup153 (top, blue; PDB 4U0C) and CPSF6 (bottom, purple; PDB 4U0B) peptides bound. Buried FG motifs are shown in orange, with key CA residues in green. **b-g**, Binding of Nups to wildtype (WT) or mutant CLPs measured by FFS. **h**, Domain architectures of Nup214, Pom121 and Nup98. FG motifs are shown as black bars. TM: Transmembrane domain, NE: inner nuclear envelope-binding region, NLS: nuclear localisation signal, GLEBS: Gle2-binding sequence, APD: Autoproteolytic domain. Domain boundaries of truncation constructs are shown in yellow. **i**, Binding of truncated constructs of Nup214, Pom121 and Nup98 to CLPs as measured by FFS. FG-repeat domains shown in red. **j**, Binding of Nup98-peptides and fragments of Nup98 to CLPs. Error bars show standard deviation.

To investigate how the capsid recognises FG-repeat domains, we chose four of our strongest binders (Nup42, Nup98, Pom121 and Nup214) and examined CA mutant binding to Nup truncations. These FG-Nups demonstrated similar profiles to the Nup153-peptide (Fig 2d-g, S2-2). In each case binding was significantly reduced by N57D, but unaffected (and possibly even enhanced) by N74D or A77V. The results obtained with these CLP mutations indicate that the FG-Nups interact with CA N57 in the FG-binding pocket, and that the binding footprints more closely resemble that of Nup153_1407-1423_ than CPSF6_313-327_. These observations may explain why N74D and A77V (CPSF6 binding mutants) retain the ability to infect non-dividing cells^27, 33–35^, as they maintain FG-interaction and use of the NPC, whereas N57 mutants (FG binding mutants) are dependent on cell division for maximal infectivity^27, 35^.

Nup98, Pom121, and Nup214 have distinct domain architectures, with each possessing a clearly defined FG-repeat domain with 44, 24 and 40 FG-motifs respectively (Fig. 2h). To determine whether the FG-domains are solely responsible for CA binding, we performed side-by-side FFS measurements comparing their FG- and non-FG-domains. Strikingly, binding was observed for all three FG-domains, while there was no detectable interaction for any non-FG domain (Fig 2i, S2-3). To probe whether binding is driven predominantly by specific FG-containing motifs (as is the case for Nup153^28^), we chose the strongest binder, Nup98, and further dissected the FG-domain into three regions, each containing 13 FGs. Each of the three FG-repeat fragments bound robustly to the WT capsid (Fig 2j and S2-3), but not N57D. CLP interaction with representative Nup98 peptides containing only a single FG motif (representing both GLFG and FGFG types) was below the detection limit for FFS (Fig 2j), suggesting that capsid binding to Nup98 is driven by weak, but FG-specific, interactions enhanced by avidity.

### HIV CA displays karyopherin-like properties

Direct interaction between CA and the FG-Nups suggests that the HIV capsid can engage with the diffusion barrier of the NPC. Furthermore, these interactions are reminiscent of the low affinity, multivalent FG-binding exhibited by karyopherins which are uniquely capable of crossing the FG-rich diffusion barrier along with their bound cargoes.

A key advance in the understanding of karyopherin-mediated transport came with the observation that isolated FG-Nups spontaneously form liquid-liquid phase-separated condensates which retain the selective properties of the NPC’s diffusion barrier^31, 36^. While seminal for the mechanistic understanding of nucleocytoplasmic transport, these FG-Nup condensates have not previously been used to study viral nuclear entry. We therefore sought to produce them to explore the interplay between the HIV capsid and the NPC diffusion barrier. Nup98 was chosen as our model system as it plays a vital role in maintaining the integrity of the NPC^37, 38^, contributes the most FGs (Fig 1e), and, among the phase-separating FG-Nups, it produces condensates which most accurately recapitulate the selectivity properties of the diffusion barrier *in vitro*^32, 39, 40^. Importantly, Nup98 was also the clearest CA binder in our FFS assay (Fig 1b,d).

We produced the unstructured FG domain of Nup98 (residues 1-499) under denaturing conditions and observed the spontaneous formation of spherical, phase separated condensates (average diameter 1.3 µm) upon shock dilution into aqueous buffer (see methods for details). We tested these condensates for their selective properties by mixing them with mCherry fused to the Importin-beta-binding domain of Importin-alpha (IBB-mCherry). IBB-mCherry (33 kDa) was excluded from the Nup98 phases (Fig 3a and S3-1) but was transported into the condensates within minutes of addition of Importin-beta (Fig 3b and S3-1), thereby demonstrating the NPC-like selective properties of our Nup98 condensates. Furthermore, the addition of Importin-beta did not result in uptake of WT mCherry (27 kDa; Fig S3-2), showing that the presence of Importin-beta does not affect the size-selective properties of the Nup98 condensates. To test whether CA could enter the FG-Nup phase we fused mCherry to the C-terminus of CA (CA-mCherry, 52 kDa). At the concentrations used in our assay (45 µM) we expected CA-mCherry to exist in equal amounts as a monomer and dimer (*K_D_* ∼ 40 µM)^41^, with the latter carrying two FG binding sites. Strikingly, CA-mCherry rapidly partitioned into the Nup98 condensates (Fig 3c and S3-1) while diffusion into the condensate was almost entirely abolished upon N57A mutation (Fig 3d, g and S3-1). Next, we produced a cysteine cross-linked hexamer (CA_hexamer_-mCherry)^42^ carrying an average of one CA-mCherry per hexamer. At 182 kDa, this construct is well beyond the size limit for passive diffusion across the NPC but presents six FG-binding sites appropriately positioned relative to their neighbouring CA protomers. Again, we observed CA_hexamer_-mCherry partitioning into the condensates (Fig 3e and S3-1), an affect that was again significantly reduced by N57A (Fig 3f,g and S3-1).

**Fig. 3:**
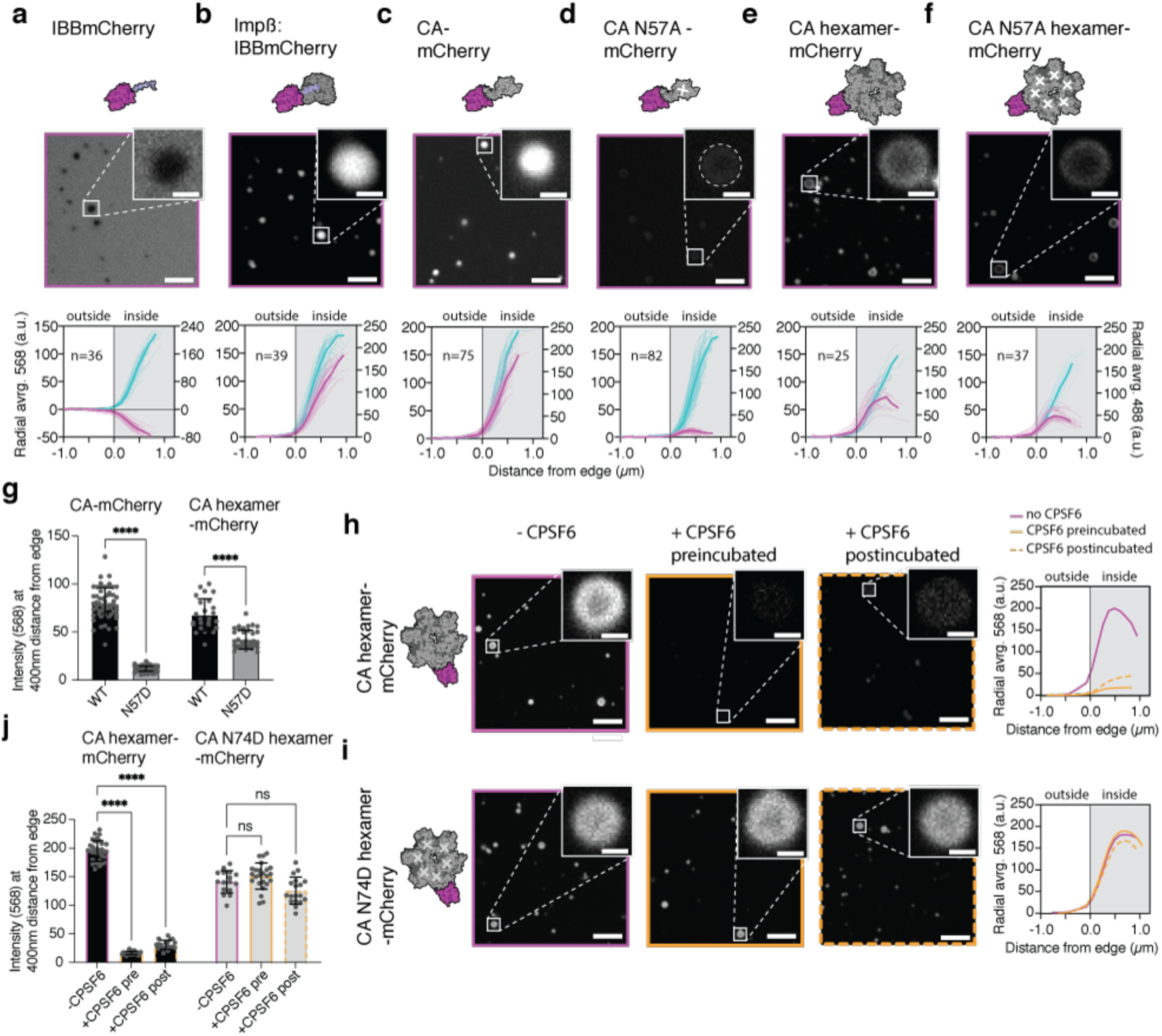
The FG-binding pocket mediates HIV-1 CA entry into Nup98 condensates. **a-f**, Single z-plane images (568 nm) of mCherry labelled controls or CA-constructs (top) and background-subtracted radially averaged fluorescence intensity condensate depth profiles (bottom; cyan, Nup98 channel; magenta, protein-mCherry channel). Mean intensity curves are shown in bold. **g,** Intensity values (568 nm) for CA-mCherry and CA_hexamer_-mCherry WT and N57A at 400 nm from the condensate edge. p<0.0001. **h-i,** Single z-plane images (568 nm) for CA_hexamer_-mCherry and CA N74D_hexamer_-mCherry in the absence of CPSF6 (magenta), preincubated with CPSF6 (orange), or post-incubated with CPSF6 (dashed orange). Mean fluorescence intensity depth profiles are shown for all conditions. **j,** Intensity values for CA_hexamer_-mCherry and CA N74D_hexamer_-mCherry in absence and presence of CPSF6 at 400 nm from the condensate edge. p<0.0001. All images: scale bar for main = 5 µm, inset = 1 µm.

To further probe the functional relevance of the FG-binding pocket, we sought to test whether known cofactors could compete for CA-binding and therefore influence Nup penetration. The recently developed drug, Lenacapavir binds the FG-pocket, but was not amenable to study in our systems as it resulted in CA clustering, consistent with its known over-assembly effect^43^. As an alternative, we employed the host cofactor, CPSF6, which had no such confounding effects. CPSF6_313-327_ peptide preferentially binds to CA_hexamer_ (*K_D_* = 50 µM) over unassembled CA (*K_D_* ∼ 700 µM)^28^. As expected, pre-treatment with 500 µM CPSF6_313-327_ had no observable effect on CA-mCherry (Fig S3-3), while abolishing the ability of CA_hexamer_-mCherry to partition into the Nup98 condensates (Fig. 3h, j and S3-4). This effect was specific to the interaction with the FG-pocket because CPSF6_313-327_ had no effect on CA(N74D)_hexamer_-mCherry which does not bind CPSF6_313-327_ (Fig. 3 i,j and S3-5). Post-treatment with CPSF6_313-327_ also removed CA_hexamer_-mCherry from the Nup98 condensates. The effects of CPSF6 not only support the role of the FG-binding pocket in mediating capsid entry into the NPC, but the pre- and post-treatment results indicate that competing cofactors may be able to influence both NPC engagement and disengagement. Indeed these results are consistent with the proposed ‘ratchet’ mechanism by which CPSF6 extracts the capsid from the nuclear face of the NPC^13^.

While these results indicate that CA can enter the NPC autonomously, previous studies have suggested that the karyopherins Importin α3^44^, Transportin-1 (TNPO1)^45^, and Transportin-3 (TNPO3)^46, 47^ each play a role in HIV-1 nuclear entry through a direct interaction with the capsid. However, we observed no specific binding between capsid and Importin α, TNPO1 or TNPO3 by FFS (Fig S3-6). Furthermore, rather than enhancing CA entry into Nup98 condensates, these karyopherins reduced entry (Fig S3-7, S3-8) consistent with the prediction that CA and karyopherins compete for FG-binding and therefore display a similar mechanism of action. Thus, we argue that our observation of CA penetrating FG condensates and facilitating the entry of an mCherry pseudo-cargo demonstrates that the HIV capsid protein possesses intrinsic karyopherin-like properties and can traverse the NPC independent of the canonical nucleocytoplasmic transport factors.

### HIV capsid-like particles partition into Nup98 condensates

In order to chaperone the viral genome and enzymes across the nuclear envelope, the HIV capsid must be capable of entering the NPC intact. To investigate the Nup-penetration properties of complete capsids, we induced co-assembly of CA(A204C) with CA(A204C)-mCherry at a ratio of 200:1. These fluorescent CLPs formed the expected heterogeneous mix of lattice structures that model the pleiomorphic capsids found in HIV virions (Fig S4-1). Confocal microscopy revealed that upon addition of CLPs, fluorescent puncta were recruited to the periphery of the Nup98 condensates (Fig 4a, S4-2), but could not resolve whether these structures had penetrated the condensates, nor whether the structural integrity of closed CA cones or spheres is maintained upon entry.

**Fig. 4:**
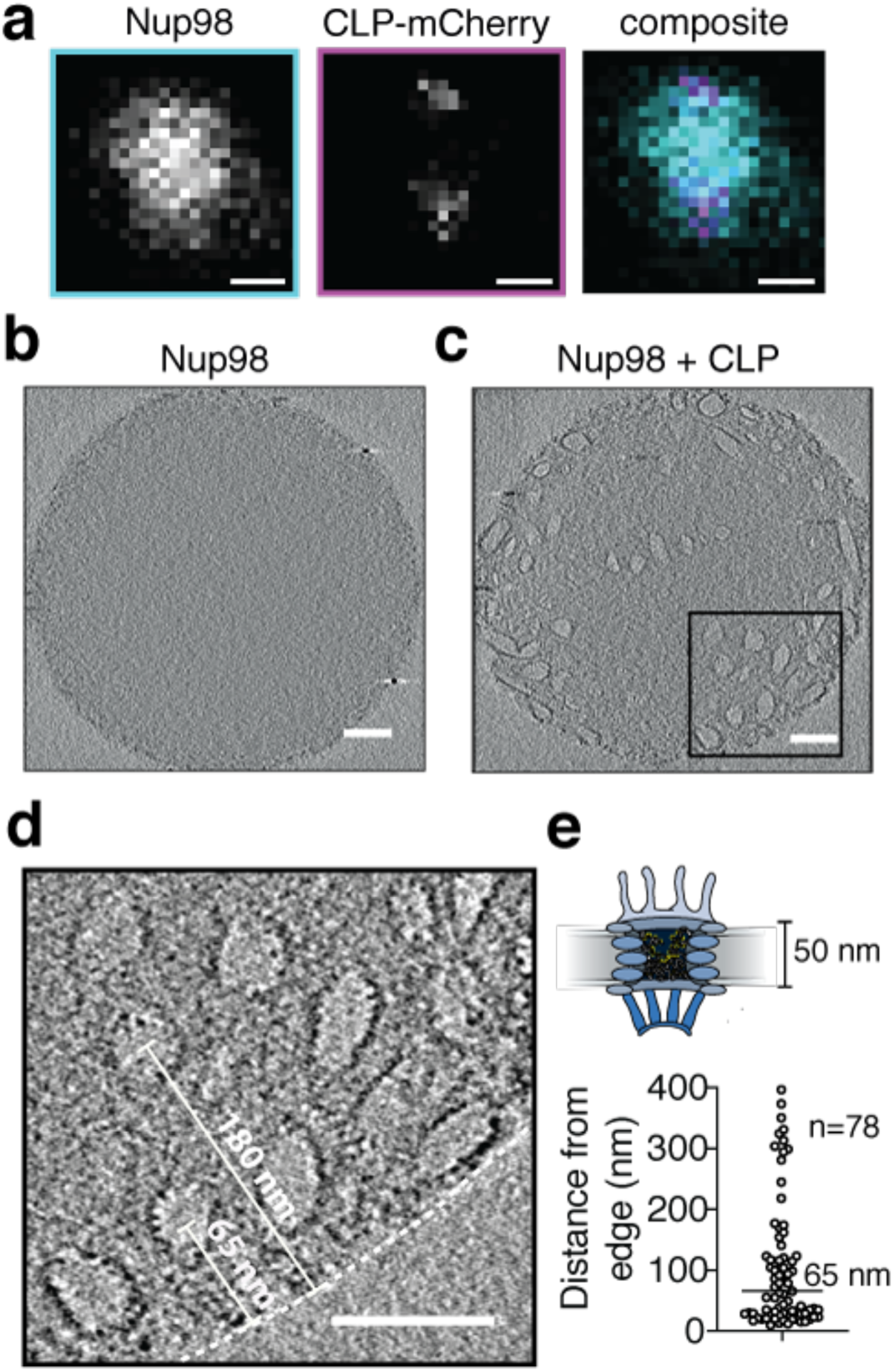
Intact HIV-1 capsid-like particles penetrate Nup98 condensates. **a,** CLP-mCherry assemblies (puncta) recruited to Nup98 condensates. Single z-plane images for 488nm (cyan), 568 nm (magenta) and composite. Scale bar = 500 nm. **b-c,** Representative slices from cryo-electron tomograms of Nup98 alone (b), or in the presence of CLPs (c). **d,** Zoomed image from c showing an average of 5 slices with a total thickness of 5.2 nm. Scale bars (b-d) = 100 nm. Penetration depths of CLPs are measured from the edge of the condensate (white dashed line) to the centre of the CLP. **e,** Penetration depths of CLPs measured from 8 condensates.

To examine whether complete HIV-1 capsids can penetrate the Nup98 condensates, we turned to cryo-electron tomography, a technique which, in contrast to previous measurements, also allowed us to study CA-Nup98 condensate interaction in a label-free context. We ensured that the condensates were electron-transparent by reducing the Nup98 concentration resulting in smaller condensates (diameter 400 nm – 1 µm) which we imaged with and without CLPs introduced immediately prior to freezing. In the absence of CA, Nup98 condensates displayed a uniform, featureless internal texture, consistent with a phase separated, intrinsically disordered protein (Fig 4b and S4-3). Strikingly, after addition of CLPs to preformed Nup98 condensates, we observed internalised pleomorphic substructures, including conical structures similar to authentic HIV capsids (Fig 4c-e and S4-3). These structures carried an electron-dense exterior with evidence of a repetitive lattice structure (Fig 4d). Internal negative space confirmed that CLPs were empty, showing that the integrity of the CLPs was not affected even when fully immersed in the FG-motif dense condensates. Furthermore, the median penetration depth of the CLPs was 65 nm (Fig 4e), which is greater than the 40 nm required to cross the central transport channel of the NPC^48^. These results show that complete HIV capsids have properties which enable them to sufficiently penetrate and enter the key material of the NPC’s diffusion barrier, while preventing the disordered Nups from infiltrating the capsid interior.

## Discussion

A key reason that the HIV:NPC interaction has remained poorly understood is that most Nups are essential for cell viability and cannot be manipulated *in cellulo* by RNAi or CRISPR/CAS. Nevertheless, over one third of nucleoporins (Nup62, Nup85, Nup88, Nup98, Nup107, Nup133, Nup153, Nup160, Nup214, Nup358 and Pom121) have previously been implicated in various aspects of the HIV lifecycle^27, 34, 35, 49–56^, but any molecular details concerning nuclear entry have remained elusive. By performing an *in vitro* screen of the CA:Nup interactions, we have circumvented the cellular challenges to show that the capsid specifically engages with the multitude of FG-motifs found within the NPC diffusion barrier. Importantly, we observe interactions with Nups located in the cytoplasmic filaments, throughout the central transport channel, and in the nuclear basket implying that capsid:Nup interactions occur at all points along the translocation pathway.

FG repeats have been classified into two main types based on the residues immediately upstream, FxFG/FG and GLFG. We observe that the HIV capsid interacts with both. The two types are thought to segregate within the NPC, with FxFG/FG repeats enriched at the cytoplasmic and nuclear peripheries, and GLFG repeats enriched in the central channel where they function as the main component of the diffusion barrier^3^. The only GLFG-containing Nup in the human NPC is Nup98, making it a crucial barrier that any foreign entity must overcome in order to enter the nucleus. It is therefore striking that Nup98 is the clearest CA-binder in our assays and that Nup98 condensates readily take up HIV capsids.

Structural studies of karyopherins have shown that they possess multiple surface pockets for binding FG-dipeptides and it is these FG interactions that enable karyopherins and their bound cargoes to selectively cross the diffusion barrier. A number of karyopherins have been shown to interact directly with FG-motifs of Nup42, Nup62, Nup98, Nup153 and Nup214^57–61^ and notably, four of these (Nup42, Nup98, Nup153, Nup214) are top CA-binders in our study.

The remarkable structural and functional similarities between CA:Nup and karyopherin:Nup interactions demonstrates that, rather than engaging with host nuclear-transport receptors, the HIV capsid has evolved to become the karyopherin for the protected HIV genome.

The concept that the HIV capsid partitions into specific liquid-liquid phases within the cell has several important implications. In such a framework, molecular weight does not represent a barrier to nuclear entry. Any assembly should be able to enter the nucleus provided that its dimensions do not exceed those of the central transport channel, and that it is not excluded from the FG-Nup phase. Notably, recent cryoET images of intact HIV-1 capsids traveling through nuclear pores revealed that the central channel can become wider than previously thought to accommodate the width of the complete capsid^13^. Combining these cryoET studies with our work presented here, the HIV capsid satisfies both criteria of appropriate size and penetration capability.

Furthermore, the collective effect of FG interactions can be recast in terms of surface energies and, in the simple case of the threefold intersection between capsid-condensate, capsid-cytosol, and condensate-cytosol interfaces, a wetting angle. This is important because, alongside the geometries of the capsid and the central transport channel, the surface energies determine the capillary forces that act on the capsid^62^, with potential to shed light on how capsids orient relative to the NPC, and possibly explain why, among the retrovirus family, only those which traverse the NPC, i.e. the lentiviruses, adopt a conical capsid morphology.

The NPC’s diffusion barrier is unlikely to be the only phase-separated compartment encountered by the HIV capsid. One of the best-characterised capsid binders, CPSF6, is known to partition into membrane-less compartments within the nucleus where it plays an essential role in targeting HIV integration^9, 63–66^. Impairment of the CA:CPSF6 interaction (through A77V, N74D mutations or CPSF6 knock-down^14, 15, 18^) results in arrest of CA at the nuclear membrane and integration at the nuclear periphery, suggesting the capsid cannot properly disengage from the NPC. Furthermore, cytoplasmic mislocalisation of CPSF6 (either through ectopic expression of CPSF6 lacking a nuclear localisation signal, or depletion of the dedicated CPSF6 karyopherin, TNPO3) prevents the HIV capsid from entering the nucleus^9, 21, 67, 68^. Our data are consistent with a model in which cytoplasmic CPSF6 competes for FG-Nup binding, thereby preventing nuclear entry; while nuclear CPSF6 helps extract the capsid out of the NPC into the nucleus and directs it towards the sites of active transcription prior to integration. It is worth noting that endogenous karyopherins similarly require assistance to disengage from the NPC. Importin-β, for example, is released from the NPC by Ran-GTP^69^. As such, nuclear CPSF6 likely performs an analogous release function in the karyopherin mimicry model of capsid-mediated nuclear entry.

Whether the HIV capsid maintains its structural integrity whilst embedded within an FG-phase (be it NPC or CPSF6) remains an open question. Furthermore, it has been proposed that the capsid ultrastructure may remodel upon transit of the NPC^13, 16, 70, 71^. While we have not explicitly addressed this question with our crosslinked capsid model, the high concentration of FG’s in the NPC (>100 mM^25^) suggests that many FG-pockets will likely be occupied simultaneously. High occupancy of the FG-pocket with the capsid targeting drug Lenacapavir or the related compound, PF74, affect capsid integrity and CA lattice stability^72^. It is therefore conceivable that effects observed only near saturation (such as changes to the capsid ultrastructure) may indeed take place during NPC transit and/or upon arrival in CPSF6-containing nuclear compartments. The role of phase partitioning in HIV uncoating has not previously been considered and will likely provide key insight into this enigmatic process.

In summary we propose a model in which the HIV capsid mimics the karyopherin mechanism of transiting the NPC by solubilising in the diffusion barrier through specific, multivalent FG-interactions. This mechanism of nuclear entry is likely conserved in other medically important viruses which may prove treatable by a strategy analogous to Lenacapavir and HIV. Certainly, further study of the HIV capsid will continue to provide a paradigm for viral infection and nuclear transport mechanisms and reveal opportunities for improving lentiviruses as gene delivery vectors along with new therapeutic approaches which may be extended to unrelated viruses.

---

**Fig. S1.**
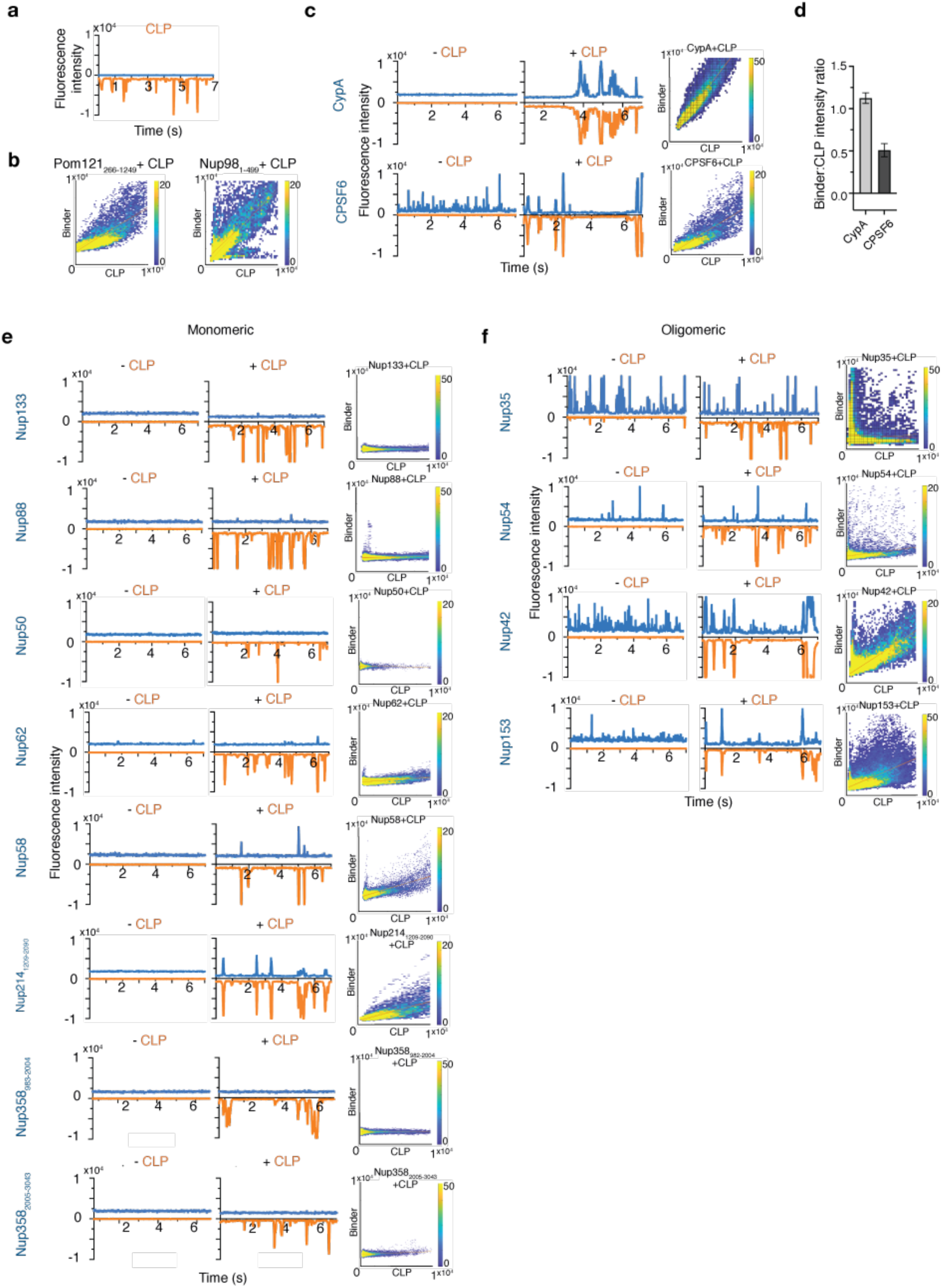
FFS data for CLPs and interactors **a**, FFS trace of AF568-CLP (orange). **b,** Heat maps of Nup to CLP fluorescence for example FFS traces shown in Fig. 1 a and b. **c, e** and **f,** FFS traces for GFP-binders in the absence and presence of AF568-CLP and heat maps of binder to CLP fluorescence intensities to calculate fluorescence intensity ratio in **d,** and Fig 1 c and d. Blue, Nup channel; orange, CLP channel.

**Table S1.** Total Number of FGs and accessible FGs calculated from AlphaFold2 RSA (relative solvent accessibility) per Nup tested in this study

| Name | Uniprot Acc | FG per Nup | Accessible FGs (AlphaFold-RSA) | Stoichiometry | Total Accessible FGs |
| --- | --- | --- | --- | --- | --- |
| Nup98 | P52948 | 40 | 40 | 48 | 1920 |
| Nup153 | P49790 | 29 | 29 | 32 | 928 |
| Nup214 | P35658 | 44<sup>^</sup> | 44 | 16 | 704 |
| Nup358* | P49792 | 22 | 19* | 32 | 608 |
| Nup58 | Q9BVL2 | 14 | 14 | 32 | 448 |
| Pom121 | Q96HA1 | 24 | 24 | 16 | 384 |
| Nup54 | Q7Z3B4 | 9 | 9 | 32 | 288 |
| Nup62 | P37198 | 6 | 6 | 48 | 288 |
| Nup42 | O15504 | 12 | 11 | 8 | 88 |
| Nup50 | Q9UKX7 | 5 | 5 | 16 | 80 |
| Nup35 | Q8NFH5 | 3 | 1 | 32 | 32 |
| Nup133 | Q8WUM0 | 3 | 1 | 32 | 32 |
| Nup88 | Q99567 | 3 | 0 | 16 | 0 |
| * | **NUP358** | **FG per Nup** | **Accessible FGs (AlphaFold-RSA)** |  |  |
|  | Nup358_0982-2004 | 11 | 11 |  |  |
|  | Nup358_2005-3043 | 8 | 8 |  |  |
|  | Nup358_3058-3224 | 3 | 0 |  |  |
|  | Total | 22 | 19 |  |  |

<sup>^</sup>Nup214 construct used in this study (1209-2090) has 42 FG
\*Due to its size, Nup358 was split into three FG-containing regions (982-2004, 2005-3043, and 3058-3224) for AlphaFold2 prediction. The total accessible FGs calculated is the sum of accessible FGs within each of these regions.

**Fig. S2-1.**
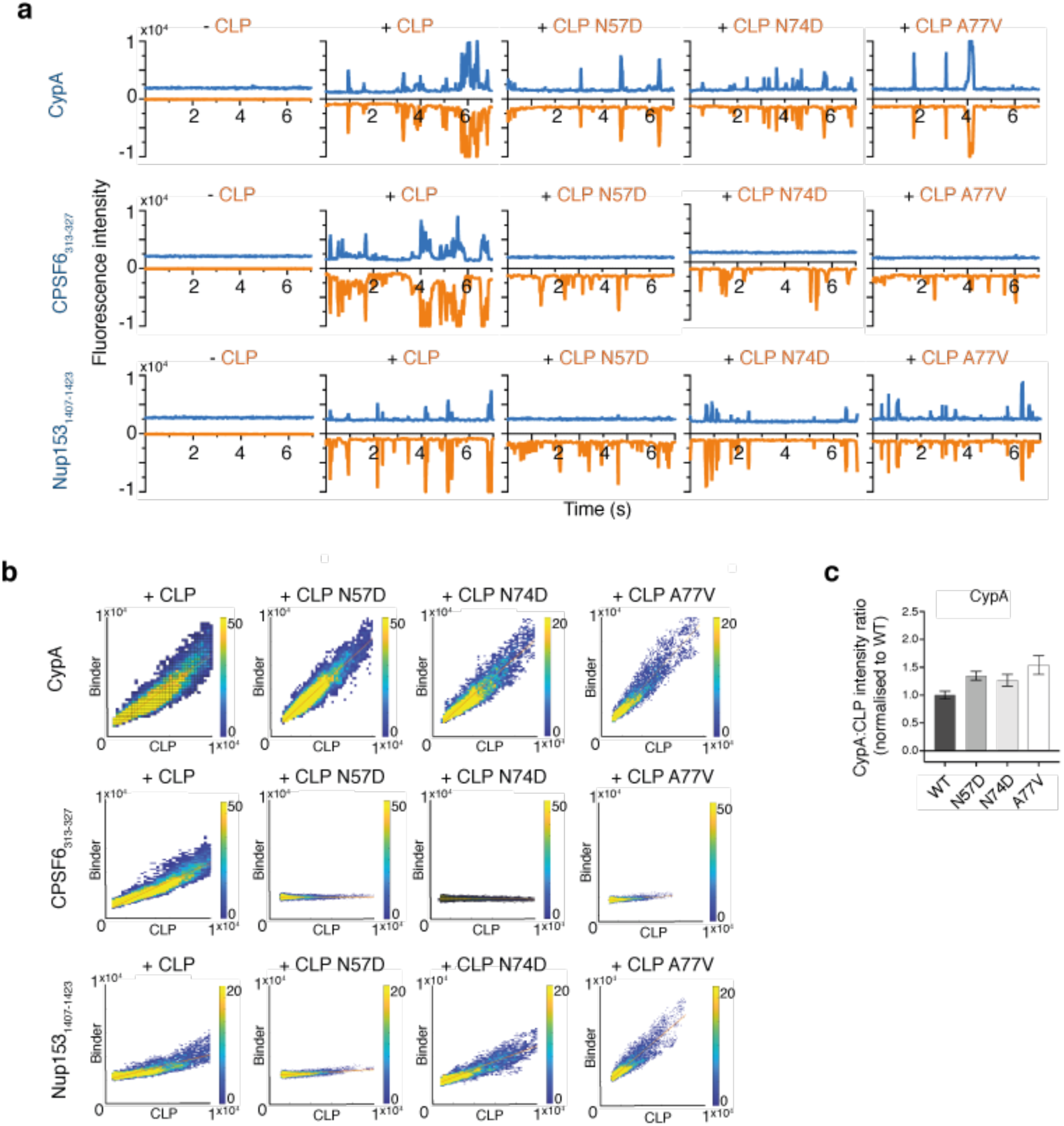
FFS data for known binders with wild-type and mutant CLPs. **a**, FFS traces for GFP-controls in the absence and presence of wild-type and mutant AF568-CLPs. Blue, Nup channel; orange, CLP channel. **b**, Corresponding heat maps of binder to CLP fluorescence to calculate fluorescence intensity ratio in **c** and Fig 2 b,c. Error bars show standard deviation.

**Fig. S2-2.**
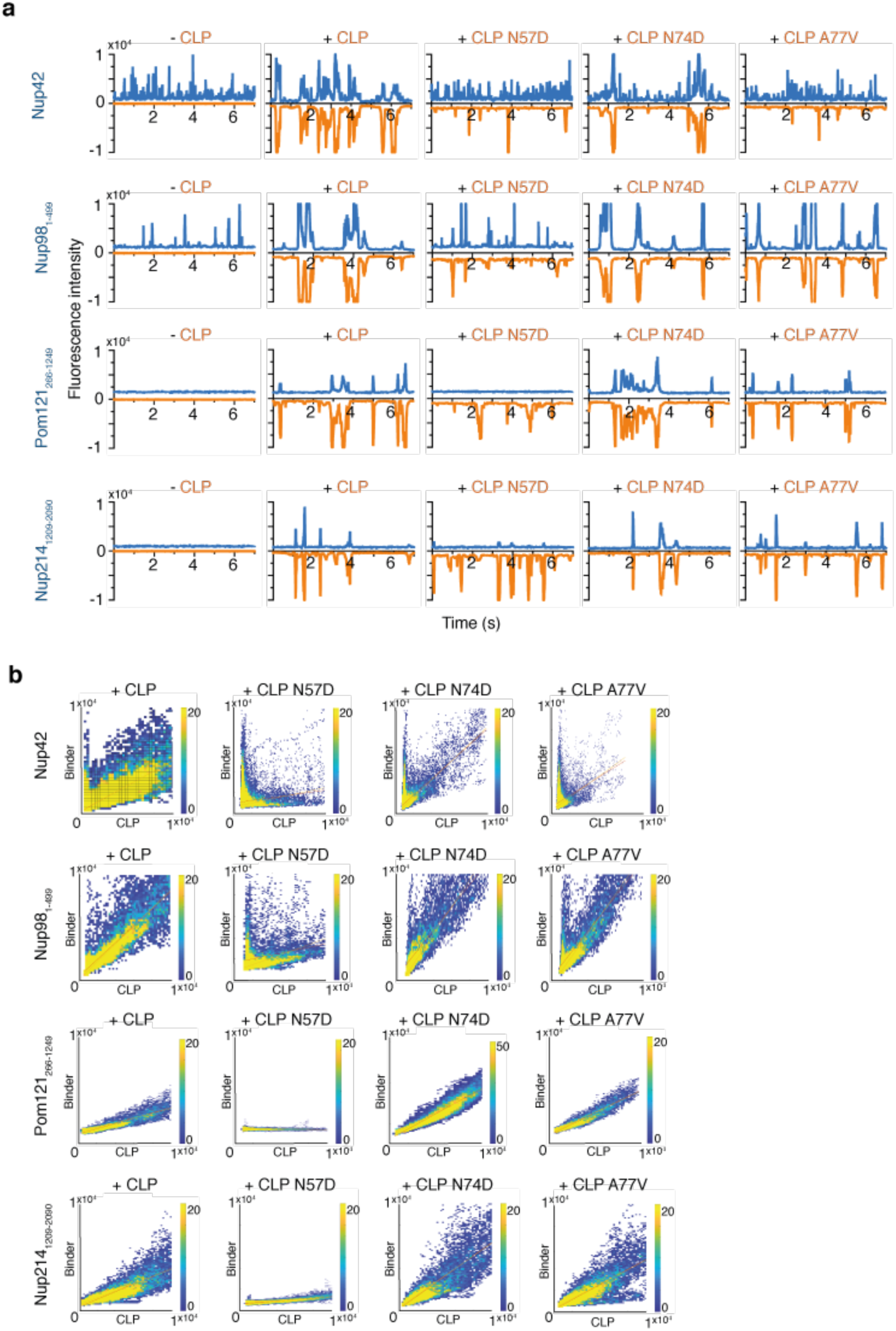
FFS data for Nup binders with wild-type and mutant CLPs. **a**, FFS traces for GFP-Nups in the absence and presence of wild-type and mutant AF568-CLPs. Blue, Nup channel; orange, CLP channel. **b**, Corresponding heat maps of binder to CLP fluorescence to calculate fluorescence intensity ratio in Fig 2 d, e, f and g.

**Fig. S2-3.**
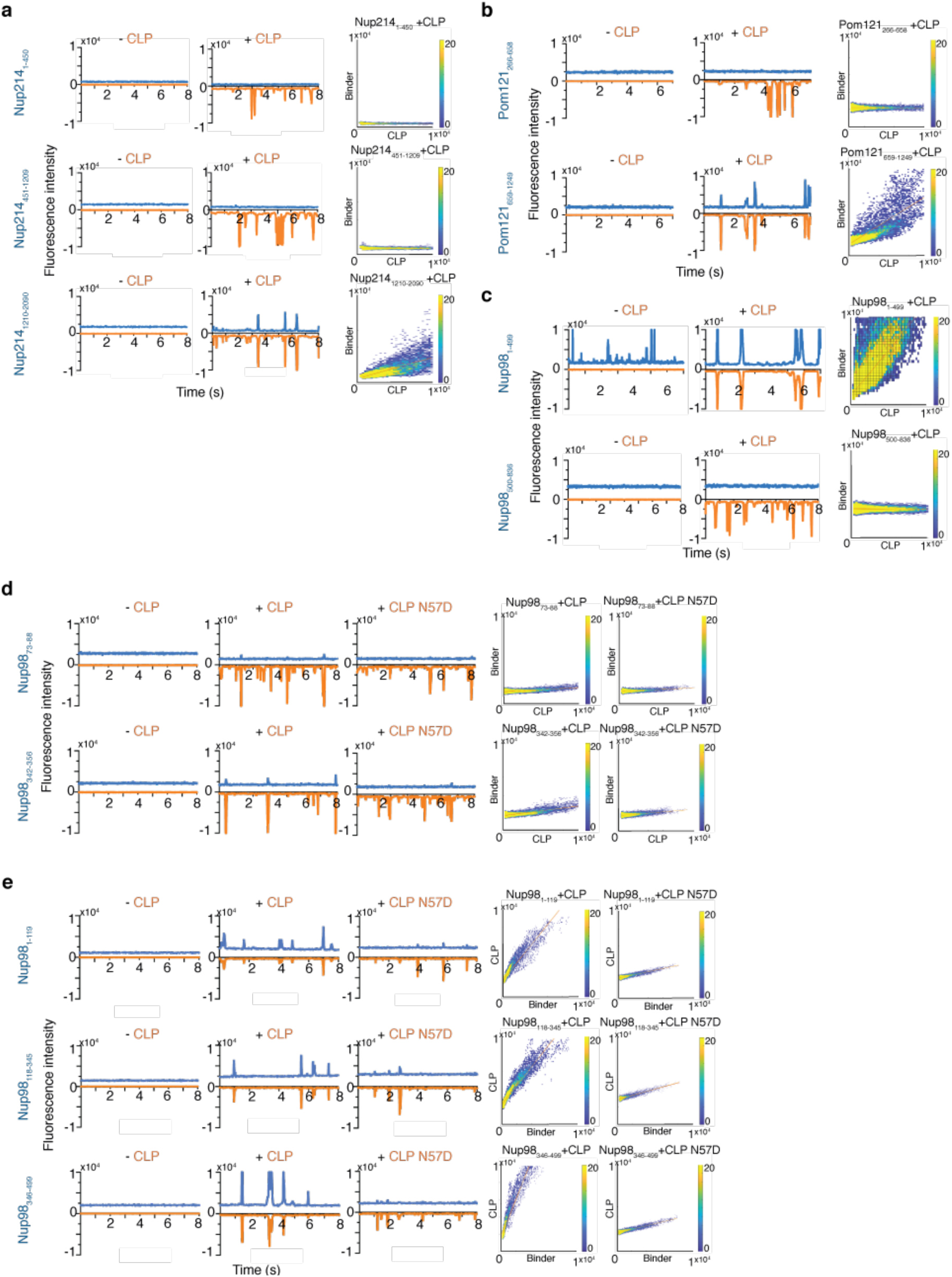
FFS data for Nup fragments with wild-type CLP and CLP N57D. FFS traces and associated coincidence heatmaps for truncation constructs of (**a**) GFP-Nup214, (**b**) GFP-Pom121, (**c**) GFP-Nup98, and (**d**) AF488-Nup98 peptides in the absence and presence of AF568-CLP (WT or N57D). **e** As a-d, with mCherry2-Nup98 fragments and AF569-CLP. For all FFS traces: blue, Nup channel; orange, CLP channel.

**Fig. S3-1:**
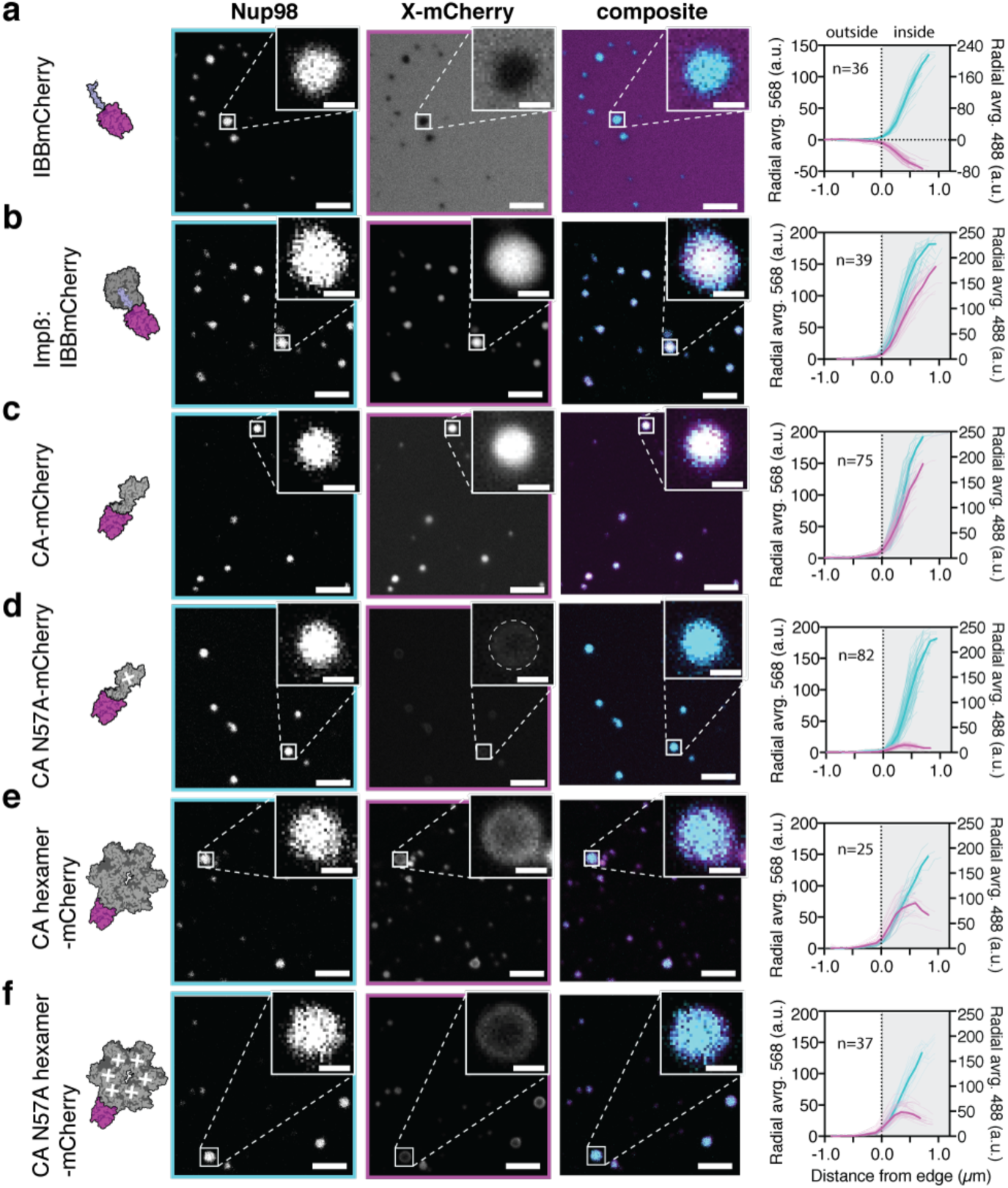
Confocal fluorescence microscopy (all channels) with Nup98 condensates and mCherry labelled controls or CA-constructs. **a-f**, Single z-plane images (cyan, 488-Nup channel; magenta, protein-mCherry channel) of (a) control fusion protein IBBmCherry (b) control complex Importin β:IBBmCherry (c) unassembled CA-mCherry (d) unassembled CA-mCherry carrying FG pocket mutation N57A (e) cross-linked CA hexamer-mCherry (f) cross-linked CA hexamer-mCherry carrying N57A. Background subtracted radially averaged fluorescence intensity across condensates for 488-Nup channel (cyan) and protein-mCherry channel (magenta). Mean intensity curves shown in bold. Scale bar 5 µm (main), 1 µm (inset).

**Fig. S3-2:**
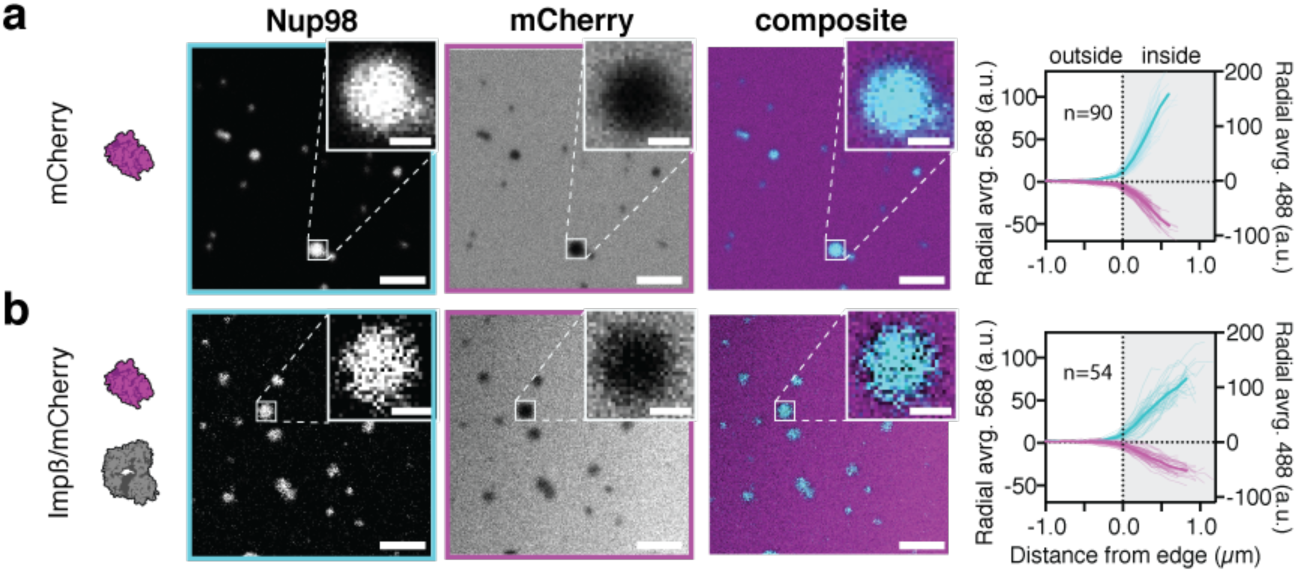
Confocal fluorescence microscopy (all channels) of Nup98 condensates with Importinβ/mCherry controls. **a-b**, Single z-plane images (cyan, 488-Nup channel; magenta, protein-mCherry channel) of (a) control mCherry (b) control Importin β and mCherry. Background subtracted radially averaged fluorescence intensity across condensates for 488-Nup channel (cyan) and protein-mCherry channel (magenta). Mean intensity curves shown in bold. Scale bar 5 µm (main), 1 µm (inset).

**Fig. S3-3:**
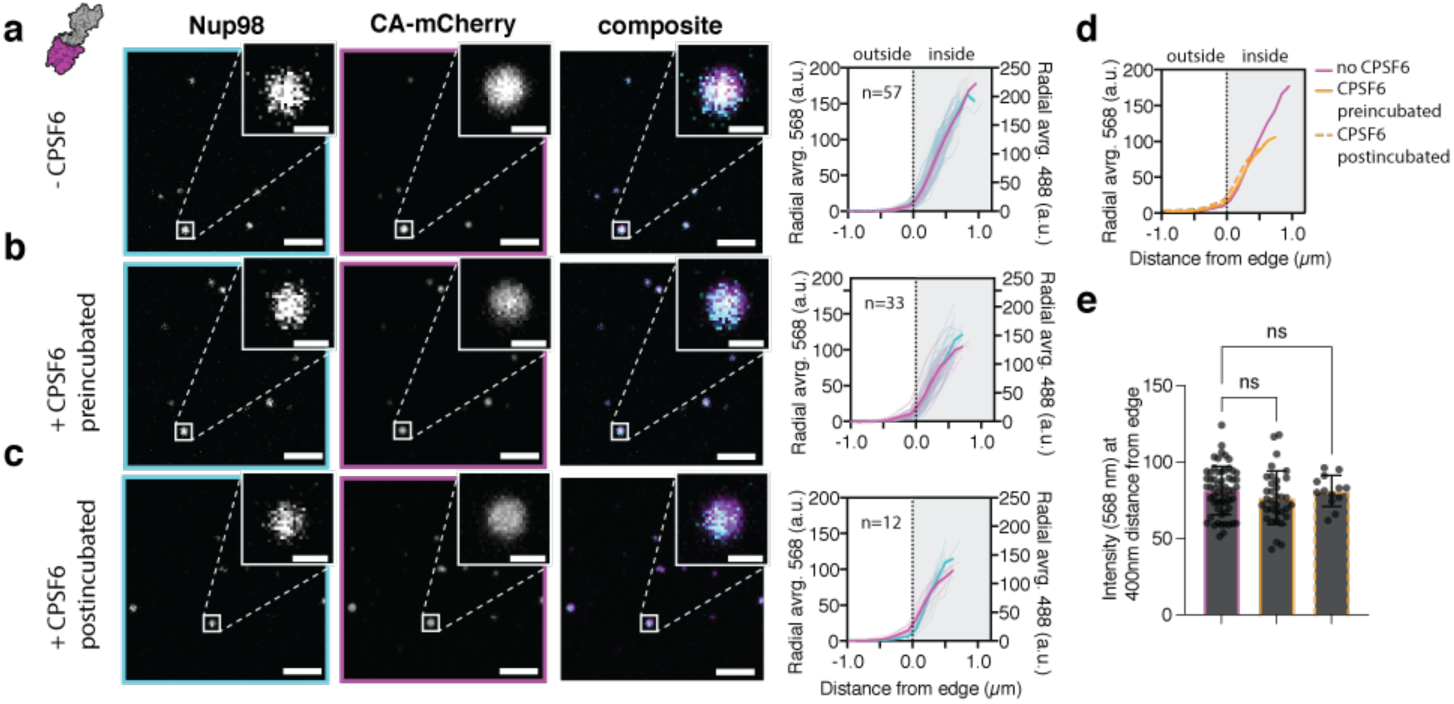
Confocal fluorescence microscopy (all channels) of Nup98 condensates and CA-mCherry in the presence of CPSF6 **a-c**, Single z-plane images (cyan, 488-Nup channel; magenta, protein-mCherry channel) of (a) CA-mCherry without CPSF6 (b) CA-mCherry with CPSF6 pre-incubated for 10 min (c) CA-mCherry with CPSF6 post-incubated 20 min. Background subtracted radially averaged fluorescence intensity across condensates for 488-Nup channel (cyan) and protein-mCherry channel (magenta). Mean intensity curves shown in bold. Scale bar 5 µm (main), 1 µm (inset). **e**, Intensity values (568) at 400 nm from edge of condensates.

**Fig. S3-4:**
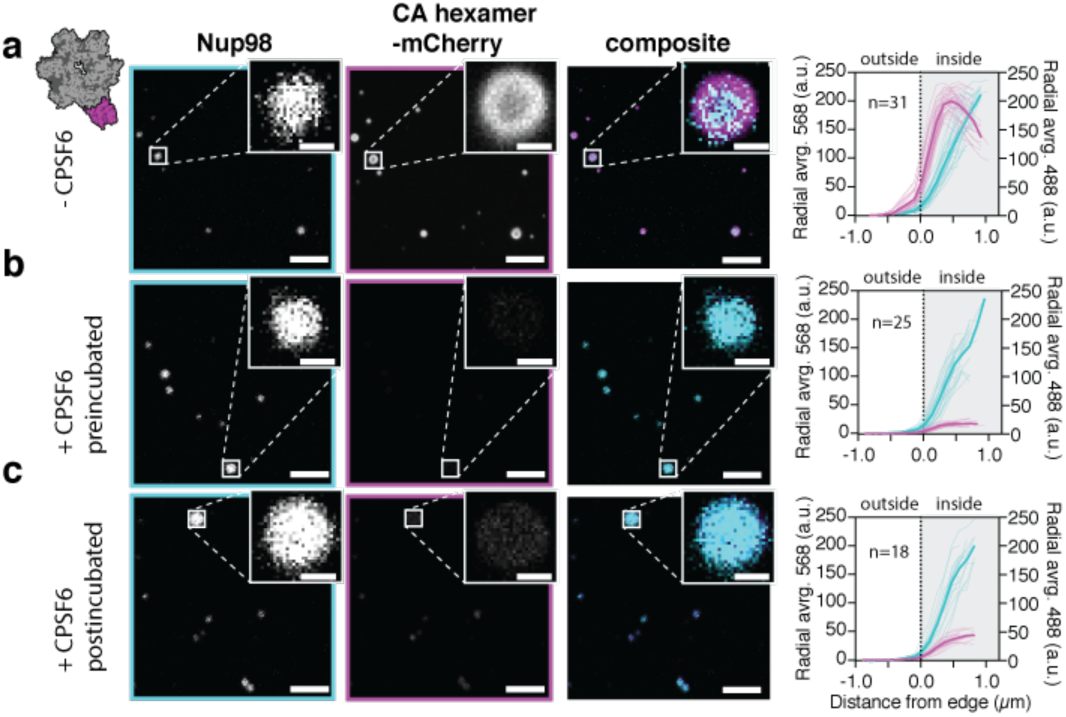
Confocal fluorescence microscopy (all channels) of Nup98 condensates and CA_hexamer_-mCherry in the presence of CPSF6 **a-c**, Single z-plane images (cyan, 488-Nup channel; magenta, protein-mCherry channel) of (a) CA_hexamer_-mCherry without CPSF6 (b) CA_hexamer_-mCherry with CPSF6 pre-incubated 10 min (c) CA hexamer-mCherry with CPSF6 post-incubated 20 min. Background subtracted radially averaged fluorescence intensity across condensates for 488-Nup channel (cyan) and protein-mCherry channel (magenta). Mean intensity curves shown in bold. Scale bar 5 µm (main), 1 µm (inset).

**Fig S3-5:**
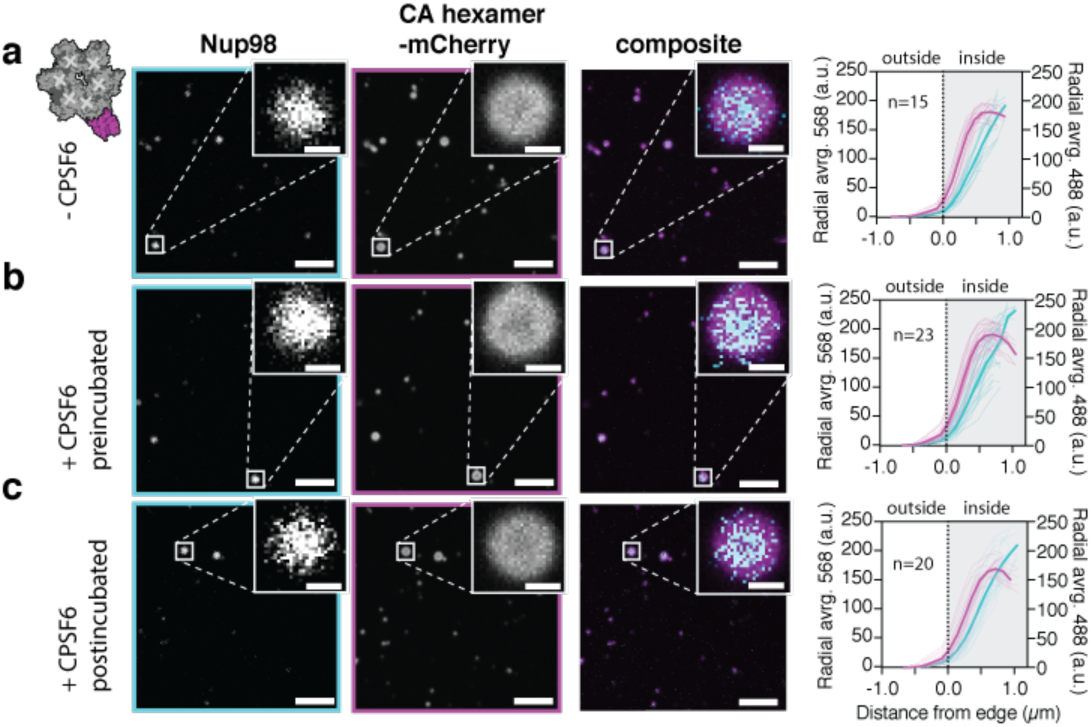
Confocal microscopy images (all channels) of Nup98 condensates and CA N74D hexamer-mCherry in the presence of CPSF6 **a-c**, Single z-plane images (cyan, 488-Nup channel; magenta, protein-mCherry channel) of (a) CA N74D hexamer-mCherry without CPSF6 (b) CA N74D hexamer-mCherry with CPSF6 pre-incubated 10 min (c) CA N74D hexamer-mCherry with CPSF6 post-incubated 20 min. Background subtracted radially averaged fluorescence intensity across condensates for 488-Nup channel (cyan) and protein-mCherry channel (magenta). Mean intensity curves shown in bold. Scale bar for main 5 µm, inset 1 µm.

**Fig S3-6:**
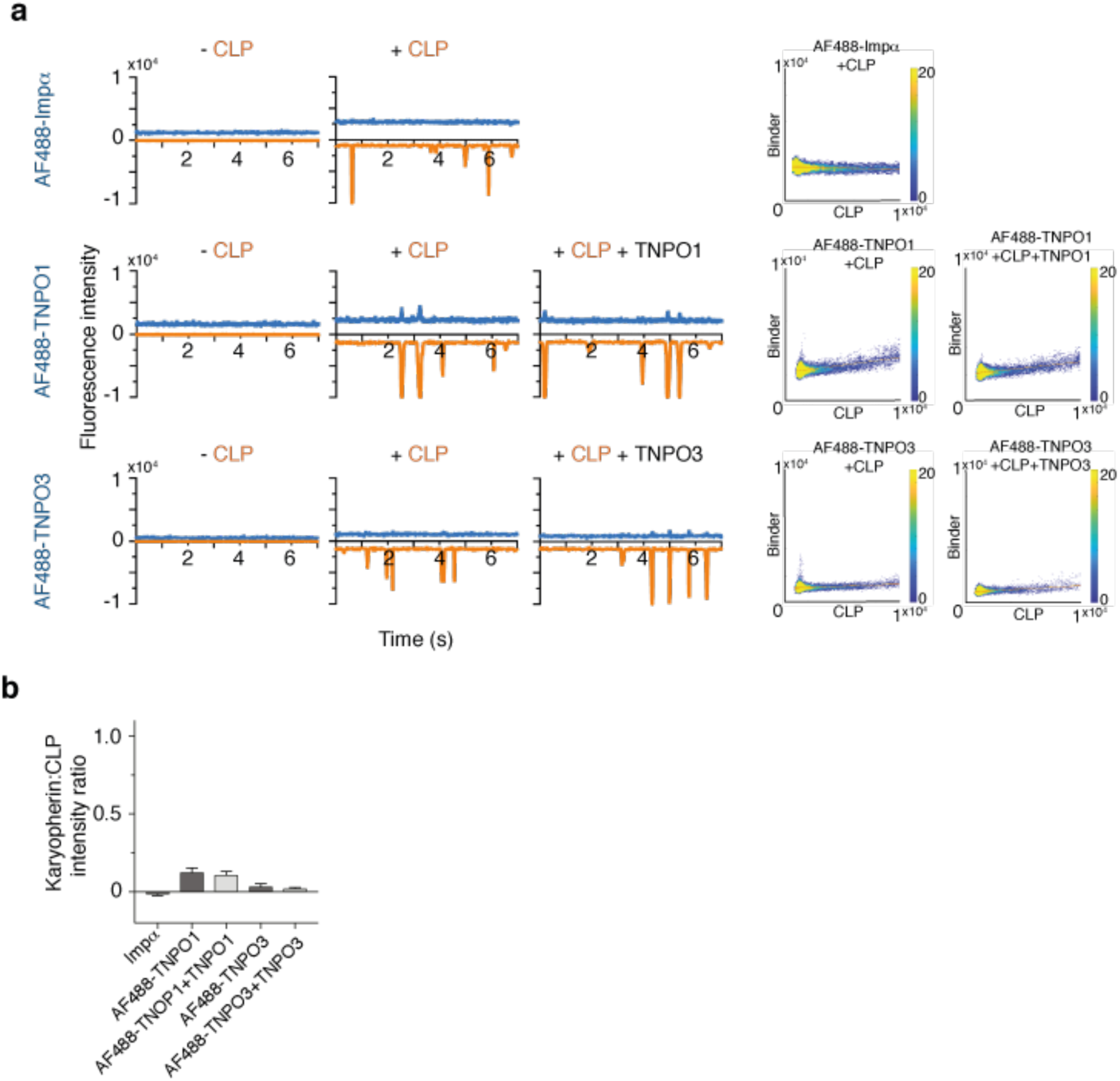
FFS shows no binding of karyopherins to CLPs. **a,** FFS traces of AF488-labelled karyopherins in absence or presence of AF568-CLP or AF568-CLP and excess unlabelled karyopherin and heat maps of karyopherin to CLP fluorescence to calculate **b**, fluorescence intensity ratio. Error bars show the standard deviation.

**Fig S3-7:**
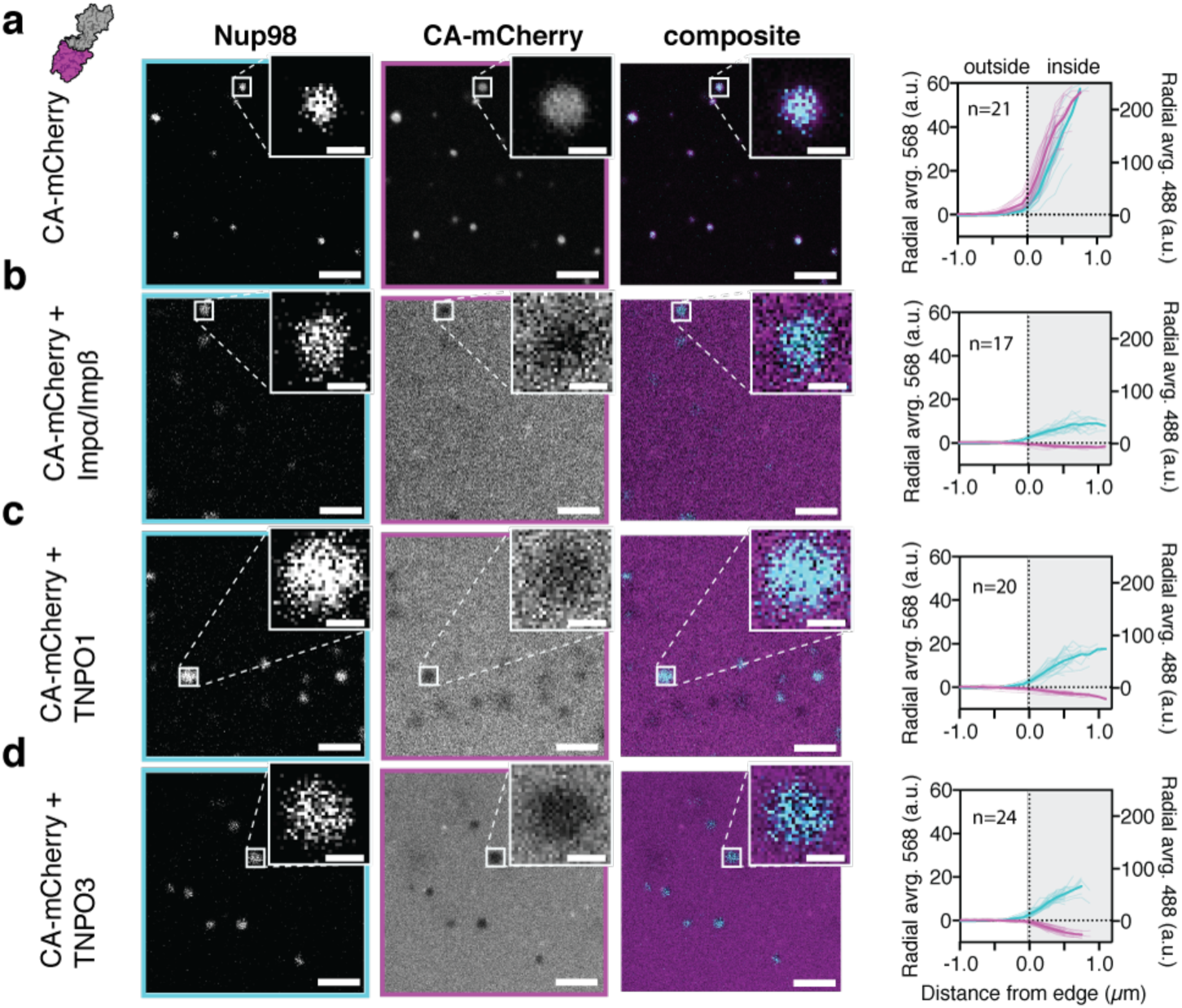
Confocal fluorescence microscopy (all channels) of Nup98 condensates and CA-mCherry in the presence of karyopherins Importinα:Importinβ, TNPO-1 and TNPO-3. **a-d**, Single z-plane images (cyan, 488-Nup channel; magenta, protein-mCherry channel) of (a) CA-mCherry (b) CA-mCherry and Importinα:Importinβ (c) CA-mCherry and TNPO-1 (d) CA-mCherry and TNPO-3. Background subtracted radially averaged fluorescence intensity across condensates for 488-Nup channel (cyan) and protein-mCherry channel (magenta). Mean intensity curves shown in bold. Nup98 signal was reduced in the presence of karyopherins, suggesting that binding of karyopherins to FG-motifs reduced the compactness of the condensates. Selective properties of the condensates were still maintained (compare Fig. S3-2). Scale bar 5 µm (main), 1 µm (inset).

**Fig. S3-8:**
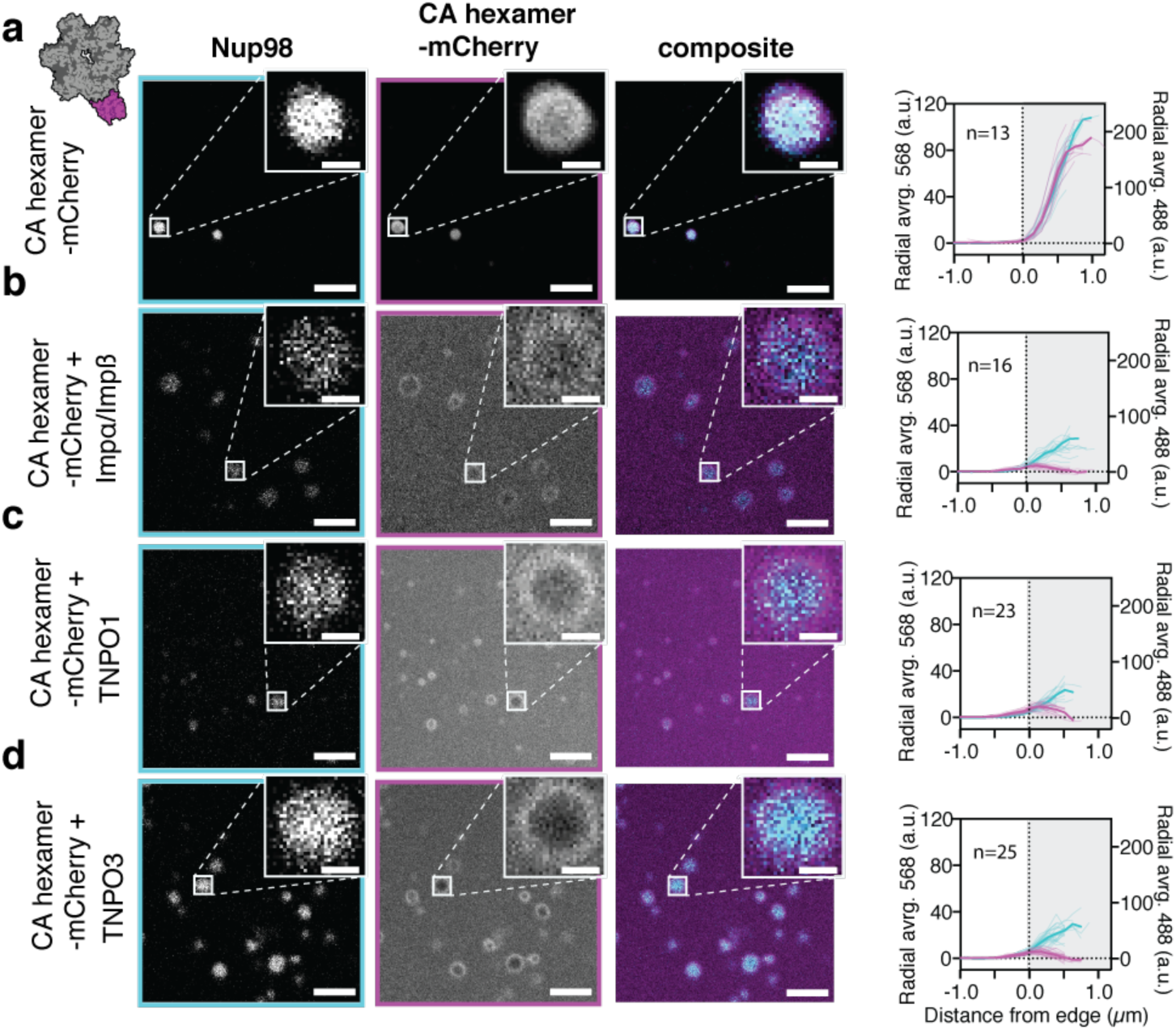
Confocal fluorescence microscopy (all channels) with Nup98 condensates and CA_hexamer_-mCherry in the presence of karyopherins Importinα:Importinβ, TNPO-1 and TNPO-3. **a-d**, Single z-plane images (cyan, 488-Nup channel; magenta, protein-mCherry channel) of (a) CA hexamer-mCherry (b) CA hexamer-mCherry and Importinα:Importinβ (c) CA hexamer-mCherry and TNPO-1 (d) CA hexamer-mCherry and TNPO-3. Background subtracted radially averaged fluorescence intensity across condensates for 488-Nup channel (cyan) and protein-mCherry channel (magenta). Mean intensity curves shown in bold. Nup98 signal was reduced in the presence of karyopherins, suggesting that binding of karyopherins to FG-motifs reduced the compactness of the condensates. Selective properties of the condensates were still maintained (compare Fig. S3-2). Scale bar 5 µm (main), 1 µm (inset).

**Fig. S4-1:**
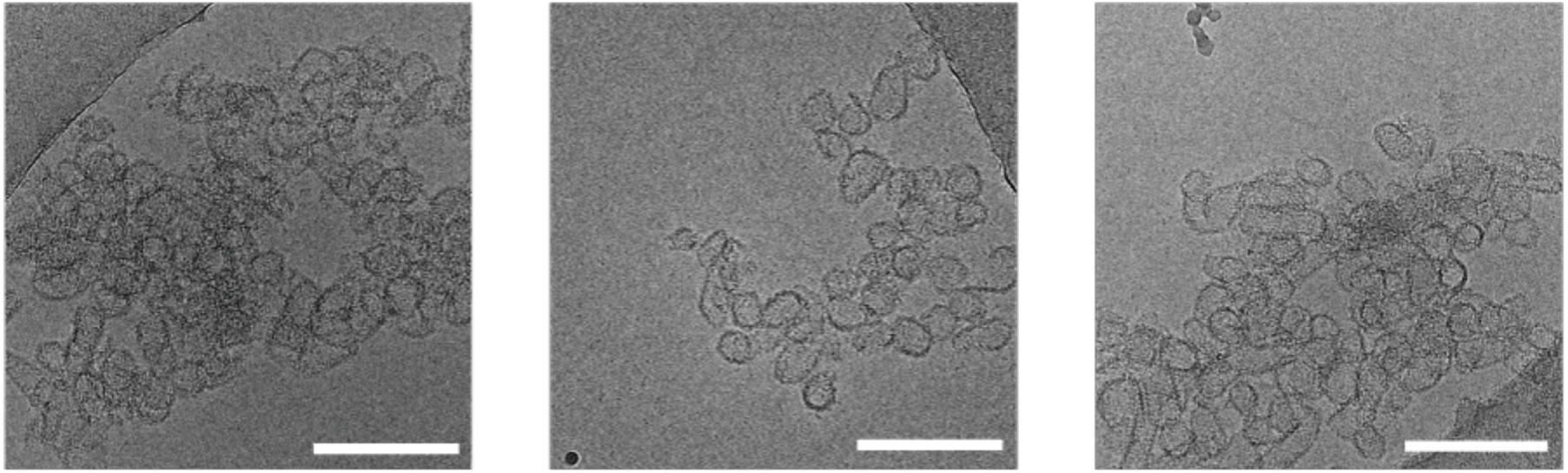
Cryo-electron microscopy images of CLP-mCherry used for confocal microscopy in Figure 4 A and S4-2. Scale bar = 200 nm.

**Fig. S4-2:**
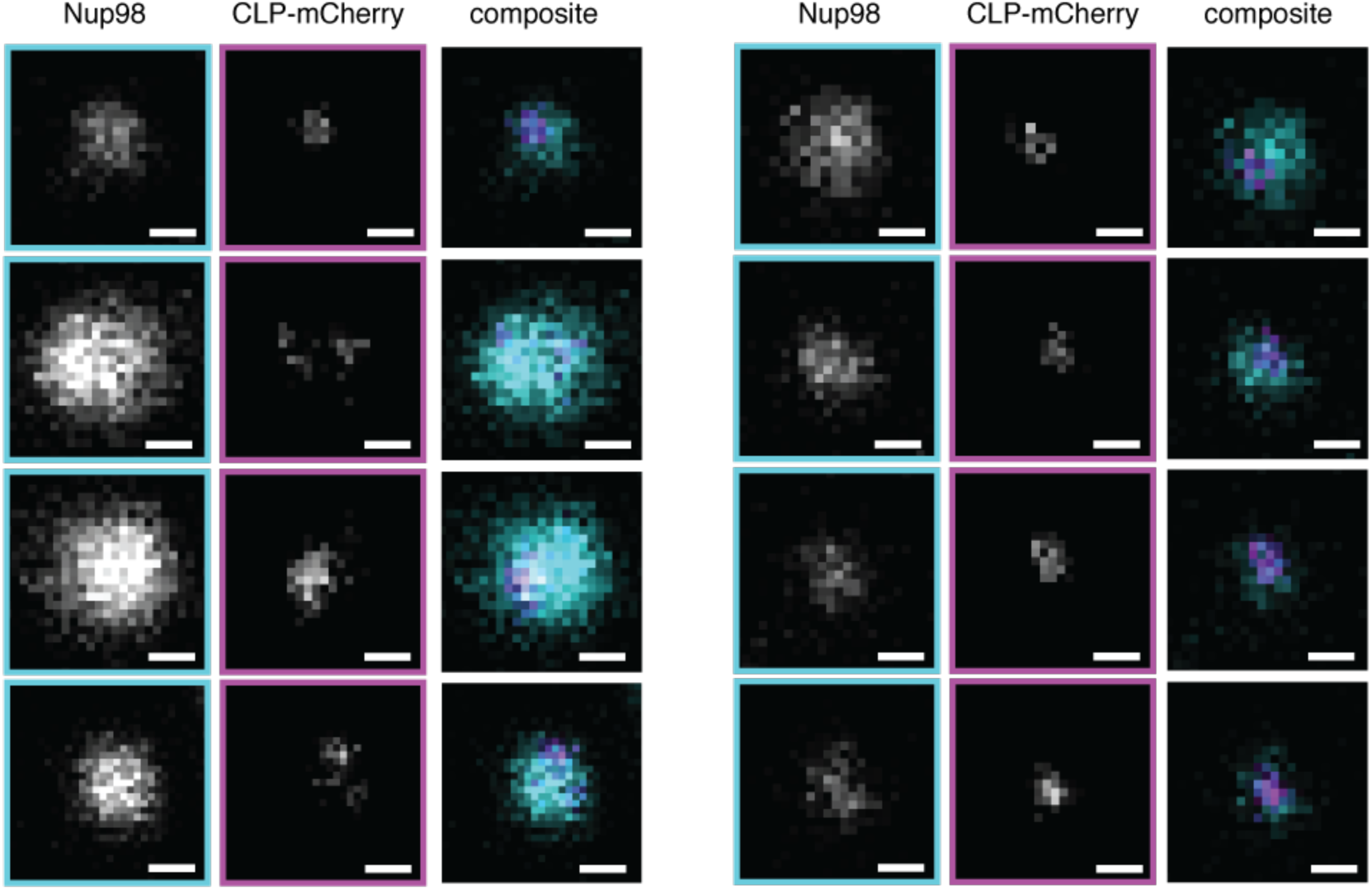
Confocal fluorescence microscopy (all channels) of CLPs bound and partitioned into Nup98 condensates CA-mCherry assemblies (puncta) recruited to Nup98 condensates. Single z-plane images for 488nm (cyan), 568 nm (magenta) and composite. Scale bar 500 nm.

**Fig. S4-3:**
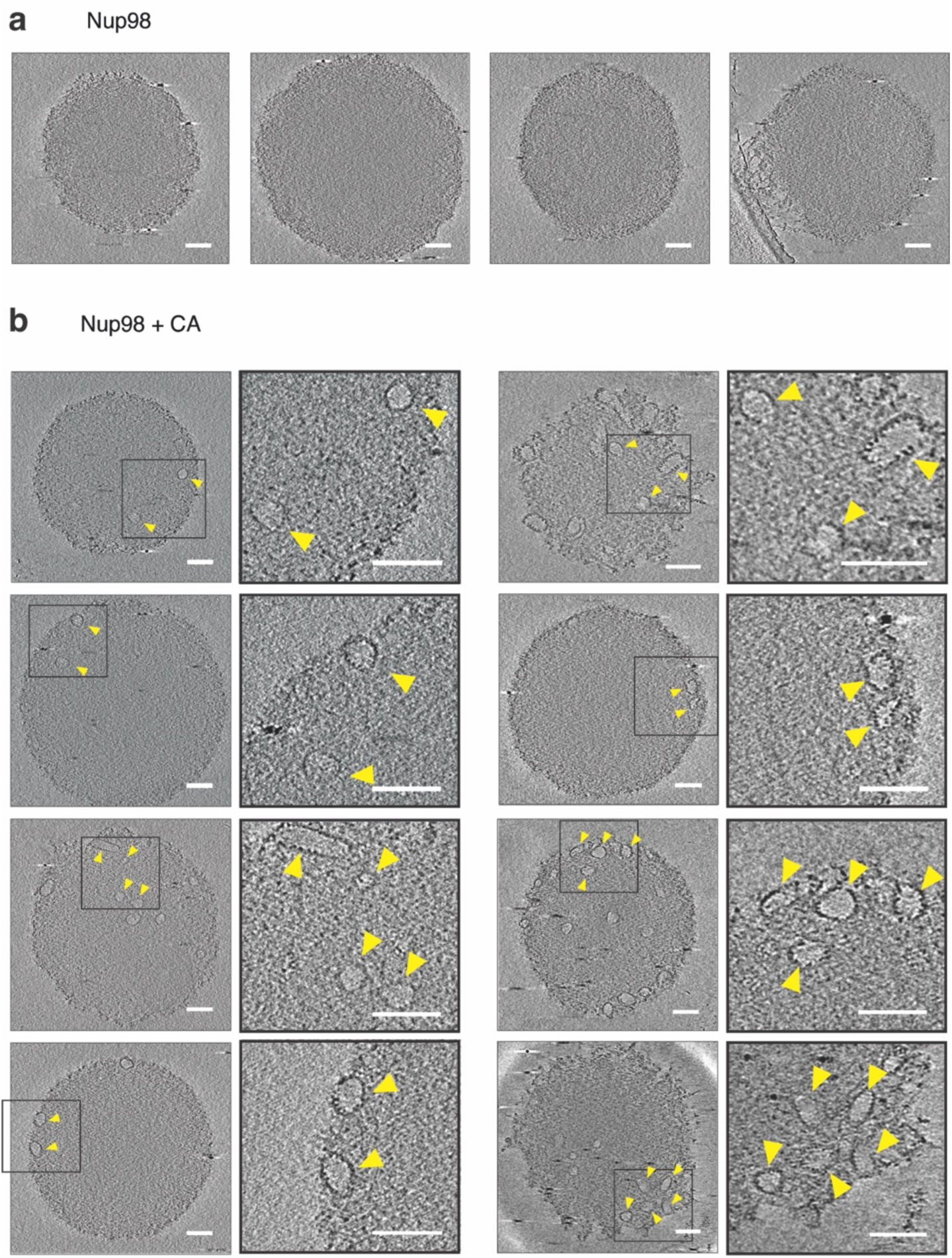
Cryo electron tomography of CLPs bound and partitioned into Nup98 condensates **a-b,** Representative slices from cryo-electron tomograms of Nup98 alone (a), or in the presence of CLPs (b). Scale bars 100 nm. Tomograms are binned by 2 and displayed as an average of 5 slices with a total thickness of 5.2 nm.

## Materials and Methods

### Constructs

Nup50, 54, 62, 88, 133, 214 or L2 in the pDONR221 gateway master vector were purchased from DNASU and cloned into the cell free expression vector pCellFree_03^1^, using the Gateway method, to produce fusion proteins with an N-terminal GFP tag (see table S2 for details). Nup35, 58, 98, 358 and Pom121 were cloned from cDNA prepared from 293T cells. Briefly, RNA was extracted from 293T cells using the Isolate II RNA mini kit (Bioline) and cDNA synthesised using oligo(dT) primers and SuperScript IV (ThermoFisher Scientific). Target genes were then cloned into pCellFree_03 using the Gibson assembly protocol (New England Biolabs). Nup98 and Pom121 fragments were purchased as gBlocks from IDT for cloning into cloned into pCellFree_03. pET28a vectors containing mCherry2-Nup98 fragments, as well as pGEX-6P-3 vectors containing Importin-*a*, TNPO1 and TNPO3 were purchased from Genscript.

### Protein expression and purification

#### HIV-1 CA proteins

HIV-1 CA proteins (K158C, A204C, R18G/A204C, N57D/A204C, N74D/A204C, N77V/A204C) were expressed in *E. coli* C41 or Rosetta2 cells and purified as previously described^2^. HIV-1 CA K158C was labelled with Alexa Fluor 568-C5-maleimide (AF568, Thermo Fisher Scientific, A20341) according to Lau et al, 2019 and mixed in an ∼ 1:200 ratio with unlabelled HIV-1 CA A204C before assembly. CA lattice assembly was carried out as described in Lau et al, 2021^3^ with the modification of a 15-minute incubation at 37 degrees and no overnight incubation.

CA_hexamer_, CA_hexamer_-mCherry and CA-mCherry were purified based on previously described protocols^4^. Briefly, CA_hexamer_, CA_hexamer_-mCherry (pOPT) and CA-mCherry (pET21a) were expressed in *E. coli* C41 cells. Cells were grown at 37 °C and protein expression was induced with 0.5 mM IPTG at OD 0.6 and cell growth was continued over night at 18 °C. Cells were harvested (4000g, 10 min, 4 °C), the cell pellets resuspended in lysis buffer (purification buffer 50 mM Tris pH 8, 50 mM NaCl, 2 mM DTT; + cOmplete EDTA-free Protease Inhibitor (Roche)) and lysed by sonication. The lysate was cleared by centrifugation (16000 g, 60 min, 4 °C) and an ammonium sulfate (20 % w/v) precipitation was carried out on the soluble fraction. After stirring for 30 min at 4 °C, the precipitated material was pelleted (16000g, 20 min, 4 °C). CA_hexamer_ was further purified by resuspending the pellet in 100 mM citric acid pH 4.5, 2 mM DTT and dialysed against the same buffer three times. The precipitated protein was pelleted (16000, 20 min, 4 °C) and the soluble fraction collected for further purification. CA_hexamer_-mCherry and CA-mCherry could not be purified with citric acid precipitation due to a possible loss of mCherry fluorescence. The ammonium sulfate precipitated pellets were resuspended in 8 ml purification buffer per litre culture and run over a 20 ml Hi-TRAP Q column in purification buffer and eluted with a gradient 1-100 % with buffer B (50 mM Tris pH 8, 500 mM NaCl). Protein containing fractions were pooled.

CA-mCherry was further purified with gel filtration (S200 in purification buffer) and fractions containing clean protein pooled and snap frozen to be stored at −80 °C.

To assemble the final cross-linked hexameric CA_hexamer_-mCherry, CA_hexamer_ and CA_hexamer_-mCherry were mixed in a ratio 5:1 and assembled via the following dialysis steps: twice against 50 mM Tris pH 8, 1000 mM NaCl, 2 mM DTT; twice against 50 mM Tris pH 8, 1000 mM NaCl; twice against 50 mM Tris pH8, 40 mM NaCl. The assembled CA_hexamer_-mCherry was finally purified with gel filtration (S200) and fractions containing hexameric CA_hexamer_-mCherry carrying one mCherry were pooled and snap frozen to be stored at – 80°C.

#### Cell-free expression

Cell free expression of GFP fusion proteins was performed as in Lau et al, 2021^3^.

#### Importinb, IBB-mCherry, Nup98

The mCherry protein was kindly provided by Jesse Goyette from the Goyette group. Importinβ and IBB-mCherry in pET11a (Genscript) were expressed as His-fusion proteins in *E. coli* Rosetta2 cells. Cells were grown at 37 °C and protein expression was induced with 0.5 mM IPTG at OD 0.6 and cell growth was continued over night at 18 °C. Cells were harvested (4000g, 10 min, 4 °C), the cell pellets resuspended in lysis buffer (purification buffer: 20 mM Tris pH 7.5, 150 mM NaCl, 2 mM DTT; + cOmplete EDTA-free Protease Inhibitor (Roche)) and lysed by sonication. The lysate was cleared by centrifugation (16000 g, 45 min, 4 °C) and the supernatant was bound to Ni-beads (Nickel sepharose high performance, Cytiva) preincubated in purification buffer, imidazole added to the protein-Ni slurry to a final concentration of 10 mM and incubated for 2 hours at 4 °C. The beads were washed with 20 CV wash buffer (purification buffer + 20 mM imidazole) and eluted in 2 ml fractions with elution buffer (purification buffer + 500 mM imidazole). Protein containing fractions were pooled and cleavage of the His-tag was achieved with TEV (IBB-mCherry) or SUMO (Importinβ) in over-night dialysis against TEV-cleavage buffer (50 mM Tris pH 7.5, 1 mM DTT) or SUMO-cleavage buffer (50 mM Tris pH 7.5, 150 mM NaCl, 1 mM DTT). Cleaved proteins were added to Ni-beads and the flow-through collected. Proteins were finally purified with SEC (S200 16/60) in purification buffer. Clean protein fractions were pooled and snap frozen to be stored at −80 °C.

Nup98 in pET28a was purified as described above with the following changes: buffers used were purification buffer (6 M GuHCl, 50 mM TrisHCl pH 7.5, 2 mM DTT), wash buffer (6 M GuHCl, 20 mM imidazole pH 8) and elution buffer (500 mM GuHCl, 500 mM imidazole pH 8). After induction with IPTG cells were grown further for 3 hours at 30 °C before harvest. Sufficient purity of the protein was achieved with affinity chromatography, consequently no SEC was performed. The His-tagged was not cleaved off. The protein-Ni slurry was stored at 4 degrees for phase separation assays.

#### Importina, TNPO1, TNPO3

Importinα, TNPO1 and TNPO3 were expressed as GST-fusion proteins from *E. coli* Rosetta2 cells, as described above for Importinb and IBBmCherry. Cell pellets were then resuspended in 25 mM Tris pH 7.5, 500 mM NaCl, 1 mM DTT, 10 % glycerol, 1x cOmplete, EDTA-free Protease Inhibitor (Roche) and lysed by sonication. Cell lysates were subject to centrifugation (18000 g, 60 min, 4 °C). Clarified lysates were bound to GSH-resin and washed with 25 mM Tris pH 7.5, 150 mM NaCl, 1 mM DTT, 10% glycerol. On column cleavage of the GST-tag was achieved with PreScission Protease. Cleaved proteins were subjected to SEC, over a Superdex 200 26/600 (Cytiva), equilibrated in 50 mM Tris pH 7.5, 200 mM NaCl, 1 mM DTT, 10% glycerol. Eluted proteins were concentrated, snap frozen and stored at – 80 °C.

### Fluorescence fluctuation spectroscopy

Fluorescence fluctuation spectroscopy was performed as described in Lau et al, 2021^3^. Briefly cell-free expressed GFP-Nup proteins (∼50 nM; equivalent to ∼ 2500 photon count), or AF488-peptides (100 nM) were mixed with AF-568-HIV-1 CA assemblies (12 uM) in 50 mM Tris, pH 8, 150 mM NaCl. Fluorescence traces were recorded for 15 s/trace in 1 ms bins using a scanning stage operated at 1 μm/s. Typically, measurements were repeated 10-15 times per sample.

### Determining the number of FGs in relative solvent accessibility (RSA)

FG-motif accessibility was determined by calculating per-residue Relative Solvent Accessibility (RSA) from AlphaFoldDB structures to determine order/disorder^5^. RSA based order/disorder for all Nups except Nup358 were accessed through Mobi-DB^6^.

Due to its length, AlphaFold-2 predictions of Nup358 were performed as three FG-containing sections (982-2004, 2005-3043, and 3058-3224) and their per residue RSA-based disorder propensity calculated locally. Binary designations of order/disorder were assigned by threshold optimised on CAID data (0.581) ^5^.

### Phase separation

For phase separation assays, the Nup98-saturated Ni slurry (see section purification Nup98. Binding capacity Ni-beads 40 mg/ml medium) was mixed with Alexa Fluor 488-C5-maleimide (AF488, Thermo Fisher Scientific) (final concentration 10 µM) and incubated for 5 min. We aimed for as little labelling with AF488 as possible with enough signal to noise to avoid confounding effects from the fluorophore for phase separation. The slurry was washed with 10 CV wash buffer 2 (500 mM GuHCl, 20 mM imidazole pH 8) + 2mM DTT, followed by a second wash step with wash buffer 2 without DTT to remove the DTT and residual fluorophore. The labelled protein was eluted with elution buffer for a final protein concentration of ∼ 4mg/ml (estimated from SDS PAGE against BSA standard) and phase separation of Nup98 was induced by a 1:10 shock dilution into assay buffer (50 mM Tris pH 7.5, 150 mM NaCl). We observed the most ‘fluid’ condensates using shock dilution, these condensates ‘hardened’ within minutes of forming, as tested with Importinb:IBBmCherry control. This resulted in limited equilibration of the condensates with some fluorescent substrates (e.g. as seen for CA_hexamer_-mCherry). Fluorescent substrates were added to the phase separated Nup98 condensates and immediately imaged.

### Confocal microscopy

Imaging was performed with Zeiss LSM 880 inverted laser scanning confocal microscope using a 63x oil immersion objective (NA=1.4) (Leica, Bensheim, Germany). The substrate-Nup98 reaction mixes were transferred into a 12 well silicone chamber (Ibidi) on 170 ± 5 µm cover slide. Z-stacks were taken around the centres of the phase separated Nup98 condensates (position with highest diameter) with sequential imaging at 488 and 568. Images were processed using ImageJ and Matlab.

Nup98 condensate-CA experiments were performed with mCherry labelled CA (CA-mCherry and CA_hexamer_-mCherry) as it was previously reported that fluorescent dyes like Alexa568 can non-specifically interact with Nup-condensates^7^.

Experimental details for Figure 3 a-f and S3-1, S3-2: Nup98 condensates were immediately mixed with the following proteins: mCherry and IBB mCherry (200 µM), Importinβ:IBBmCherry and Importinβ + mCherry (both 100 µM), CA-mCherry and CA N57A-mCherry (45 µM), CA_hexamer_-mCherry and CA N57A_hexamer_-mCherry (65 µM). The mixed Nup98-protein samples were imaged after around 5 min incubation on the coverslip.

Experimental details for Figure 3 h,i and S3-3, S3-4, S3-5: For the CPSF6_P_ preincubation sample, CA-mCherry, CA_hexamer_-mCherry and CA N74D_hexamer_-mCherry (65 µM final) were incubated with in MQW dissolved CPSF6 (500 µM final) for 10 min, followed by mixing with Nup98 condensates and imaging after around 5 min incubation on the coverslip. The control sample and the CPSF6 postincubation sample were incubated with MQW for 10 min, followed by mixing with Nup98 condensates and imaging after around 5 min incubation on the coverslip. For the CPSF6 postincubation sample, CPSF6 was added to the sample on the coverslip and imaged after 5, 10, 15 and 20 min.

Experimental details for S3-7 and S3-8: CA-mCherry and CA_hexamer_-mCherry (final 25µM) were incubated with Importinα:Importinβ, TNPO1 and TNPO3 (all final 2.5 µM), respectively, for 10 min, followed by mixing with Nup98 condensates and imaging after around 5 min incubation on the coverslip.

Experimental details for Figure 4 and S4-2: CLP-mCherry were assembled with a ratio 1:100 CA-mCherry (final 0.4 µM) to CA (final 40 µM) by addition of NaCl (final 1 mM) and a 15 min incubation at 37 °C. After assembly, the sample was spun down hard (18000g, 7 min) to separate the assemblies from non-assembled monomer. The pellet was resuspended in assay buffer (50 mM Tris pH 7.5, 150 mM NaCl, resuspension volume same as original sample volume) followed by a slow spin of the resuspended pellet (4000g, 5 min) to remove aggregates. Nup98 condensates were immediately mixed with CA-mCherry assemblies (1 µM monomer concentration) and imaged after around 5 min incubation on the coverslip.

### Radial intensity profiles

Averaged radial intensity profiles were obtained using the plugin Radial Profile in ImageJ. Z-slices were chosen at the centres of the phase separated Nup98 condensates (position with largest diameter). Profiles were background subtracted. The edge of a condensate was defined from the 488 channel as the first intensity value bigger than 5. 200 nm was subtracted from this value to account for point spread function of the fluorescent pixel. This point was defined as point 0 µm in the radial averaged intensity graphs (Figures 3, S3-1, S3-2, S3-3, S3-4, S3-5, S3-7, and S3-8).

### Cryo-Electron microscopy

Nup98 condensates were prepared as for confocal microscopy with the following modifications. To induce the formation of smaller condensates the unlabelled Nup98 was diluted 2 fold in 500 mM GuHCl, 500 mM imidazole pH 8 before a 1:10 shock dilution into assay buffer (50 mM Tris pH 7.5, 150 mM NaCl).

Unlabelled CLPs were prepared as described above. A two-step centrifugation protocol was performed as for confocal microscopy. Equal volumes of CLP suspension were mixed with freshly shock diluted condensates, immediately prior to plunge freezing to minimise sample aggregation.

#### Frozen-hydrated sample preparation

4.5 µL of freshly mixed Nup98 condensate and capsid mixture with protein A-gold (10 nm) were applied onto a glow-discharged Quantifoil R2/2 copper grid (Quantifoil Micro Tools). The grid was blotted in the front for 2.5 s at 15 °C with 90% relative humidity and then plunged into liquid ethane using a Lecia EM GP device (Lecia Microsystem). The vitrified grids were then stored in liquid nitrogen prior to cryo-ET imaging.

#### Cryo-ET imaging and reconstruction

The grids were imaged on a Talos Arctica electron microscope (Thermo Fisher Scientific) operated at 200 kV acceleration voltage. Cryo-ET data were collected with single-axis tilt on a Falcon III direct electron detector (Thermo Fisher Scientific) in linear mode at a magnification of 28,000x with a pixel size of 5.23 Å.

Tilt series were collected using the dose-symmetric scheme^8^ from −60 to 60° at 3° intervals using Tomography software (Thermo Fisher Scientific) with the defocus value set at −10 µm. Total dose for each tilt series ranged from 60-70 e/Å2. Images of tilt series were binned twofold before tomograms were reconstructed. Three-dimensional reconstructions from tilt series were generated with the IMOD package^9^. Fiducial tracking was used to align the stack of tilted images.

## Code Availability Statement

Fluorescence fluctuation spectroscopy traces were analyzed using custom software TRISTAN freely available on https://github.com/lilbutsa/Tristan.

## Acknowledgements

We thank Jeffrey Stear and James Walsh for critical reading of this manuscript. This work was supported by a National Health and Medical Research Council Ideas Grant (GNT2013215, D.A.J., T.B.) and Wellcome Trust Collaborator Award (214344/Z/18/Z, D.A.J., T.B., G.J.T.). C.F.D. was supported by a NHMRC Early Career Fellowship (GNT1110116). D.A.J. was supported by a UNSW Scientia Fellowship. The confocal imaging component of this study was carried out using instruments situated in, and maintained by, the Katharina Gaus Light Microscopy Facility at UNSW. We acknowledge the use of the Cryo Electron Microscopy Facility through the Victor Chang Cardiac Research Institute Innovation Centre, funded by the NSW government, and the Electron Microscope Unit at UNSW Sydney. We also acknowledge the use of the Recombinant Products Facility and facilities in the Structural Biology Facility within the Mark Wainwright Analytical Centre– UNSW, funded in part by the Australian Research Council Linkage Infrastructure, Equipment and Facilities Grant: ARC LIEF 190100165.

## Author contributions

C.F.D. and S.H. contributed equally to this work. C.F.D. was responsible for design and execution of fluorescence fluctuation experiments. S.H. was responsible for design and execution of nucleoporin phase separation and confocal microscopy experiments. C.F.D, S.H., J.R, and N.A. acquired and analysed cryoEM data. A.T. and N.L. contributed useful discussions around experimental design and contributed to ffs and confocal measurements. S.E and Y.G. contributed to ffs experimental design, cell-free protein expression, and data collection. S.C.A-I. and R.G.M. contributed to condensate experimental design and data interpretation. G.J.T. and T.B. contributed to experimental design and data interpretation. D.A.J. supervised all aspects of the project. C.F.D, S.H., and D.A.J. wrote the manuscript with input from all authors.

## Author information

The authors declare no competing financial interests. Correspondence and requests for materials should be addressed to D.A.J.

## References

1. Rabut, G., Doye, V. & Ellenberg, J. Mapping the dynamic organization of the nuclear pore complex inside single living cells. Nat Cell Biol 6, 1114–1121 (2004).

2. Ori, A. et al. Cell type-specific nuclear pores: a case in point for context-dependent stoichiometry of molecular machines. Mol Syst Biol 9, 648 (2013).

3. Kim, S. J. et al. Integrative structure and functional anatomy of a nuclear pore complex. Nature 555, 475–482 (2018).

4. Mosalaganti, S. et al. AI-based structure prediction empowers integrative structural analysis of human nuclear pores. Science 376, eabm9506 (2022).

5. Paine, P. L., Moore, L. C. & Horowitz, S. B. Nuclear envelope permeability. Nature 254, 109–114 (1975).

6. Weis, K. Nucleocytoplasmic transport: cargo trafficking across the border. Current Opinion in Cell Biology 14, 328–335 (2002).

7. Yamashita, M. & Emerman, M. Capsid is a dominant determinant of retrovirus infectivity in nondividing cells. J. Virol. 78, 5670–5678 (2004).

8. Qi, M., Yang, R. & Aiken, C. Cyclophilin A-dependent restriction of human immunodeficiency virus type 1 capsid mutants for infection of nondividing cells. J. Virol. 82, 12001–12008 (2008).

9. Schaller, T. et al. HIV-1 capsid-cyclophilin interactions determine nuclear import pathway, integration targeting and replication efficiency. PLoS Pathog 7, e1002439 (2011).

10. Lahaye, X. et al. The capsids of HIV-1 and HIV-2 determine immune detection of the viral cDNA by the innate sensor cGAS in dendritic cells. Immunity 39, 1132–1142 (2013).

11. Rasaiyaah, J. et al. HIV-1 evades innate immune recognition through specific cofactor recruitment. Nature 503, 402–405 (2013).

12. Zuliani-Alvarez, L. et al. Evasion of cGAS and TRIM5 defines pandemic HIV. Nature Microbiology 7, 1762–1776 (2022).

13. Zila, V. et al. Cone-shaped HIV-1 capsids are transported through intact nuclear pores. Cell 184, 1032–1046.e18 (2021).

14. Zila, V., Müller, T. G., Laketa, V., Müller, B. & Kräusslich, H.-G. Analysis of CA Content and CPSF6 Dependence of Early HIV-1 Replication Complexes in SupT1-R5 Cells. MBio 10, (2019).

15. Burdick, R. C. et al. HIV-1 uncoats in the nucleus near sites of integration. Proc Natl Acad Sci U S A 117, 5486–5493 (2020).

16. Li, C., Burdick, R. C., Nagashima, K., Hu, W.-S. & Pathak, V. K. HIV-1 cores retain their integrity until minutes before uncoating in the nucleus. Proc Natl Acad Sci U S A 118, (2021).

17. Müller, T. G. et al. HIV-1 uncoating by release of viral cDNA from capsid-like structures in the nucleus of infected cells. eLife 10, (2021).

18. Bejarano, D. A. et al. HIV-1 nuclear import in macrophages is regulated by CPSF6-capsid interactions at the nuclear pore complex. eLife 8, (2019).

19. Kubitscheck, U. et al. Nuclear transport of single molecules: dwell times at the nuclear pore complex. J Cell Biol 168, 233–243 (2005).

20. Yang, W. & Musser, S. M. Nuclear import time and transport efficiency depend on importin beta concentration. J Cell Biol 174, 951–961 (2006).

21. Tu, L.-C. & Musser, S. M. Single molecule studies of nucleocytoplasmic transport. Biochim Biophys Acta 1813, 1607–1618 (2011).

22. Tetenbaum-Novatt, J., Hough, L. E., Mironska, R., McKenney, A. S. & Rout, M. P. Nucleocytoplasmic transport: a role for nonspecific competition in karyopherin-nucleoporin interactions. Mol Cell Proteomics 11, 31–46 (2012).

23. Tu, L.-C., Fu, G., Zilman, A. & Musser, S. M. Large cargo transport by nuclear pores: implications for the spatial organization of FG-nucleoporins. EMBO J. 32, 3220–3230 (2013).

24. Milles, S. et al. Plasticity of an ultrafast interaction between nucleoporins and nuclear transport receptors. Cell 163, 734–745 (2015).

25. Aramburu, I. V. & Lemke, E. A. Floppy but not sloppy: Interaction mechanism of FG-nucleoporins and nuclear transport receptors. Seminars in Cell and Developmental Biology 68, 34–41 (2017).

26. Rebensburg, S. V. et al. Sec24C is an HIV-1 host dependency factor crucial for virus replication. Nature Microbiology 1–23 (2021). doi:10.1038/s41564-021-00868-1

27. Matreyek, K. A., Yücel, S. S., Li, X. & Engelman, A. Nucleoporin NUP153 phenylalanine-glycine motifs engage a common binding pocket within the HIV-1 capsid protein to mediate lentiviral infectivity. PLoS Pathog 9, e1003693 (2013).

28. Price, A. J. et al. Host Cofactors and Pharmacologic Ligands Share an Essential Interface in HIV-1 Capsid That Is Lost upon Disassembly. PLoS Pathog 10, e1004459– 17 (2014).

29. Bhattacharya, A. et al. Structural basis of HIV-1 capsid recognition by PF74 and CPSF6. Proc Natl Acad Sci U S A 111, 18625–18630 (2014).

30. Lau, D. et al. Rapid HIV-1 Capsid Interaction Screening Using Fluorescence Fluctuation Spectroscopy. Anal. Chem. 93, 3786–3793 (2021).

31. Frey, S., Richter, R. P. & Görlich, D. FG-rich repeats of nuclear pore proteins form a three-dimensional meshwork with hydrogel-like properties. Science 314, 815–817 (2006).

32. Labokha, A. A. et al. Systematic analysis of barrier-forming FG hydrogels from Xenopus nuclear pore complexes. EMBO J. 32, 204–218 (2012).

33. Lee, K. et al. Flexible use of nuclear import pathways by HIV-1. Cell Host & Microbe 7, 221–233 (2010).

34. Kane, M. et al. Nuclear pore heterogeneity influences HIV-1 infection and the antiviral activity of MX2. eLife 7, (2018).

35. Buffone, C. et al. Nup153 Unlocks the Nuclear Pore Complex for HIV-1 Nuclear Translocation in Nondividing Cells. J. Virol. 92, (2018).

36. Frey, S. & Görlich, D. A Saturated FG-Repeat Hydrogel Can Reproduce the Permeability Properties of Nuclear Pore Complexes. Cell 130, 512–523 (2007).

37. Wu, X. et al. Disruption of the FG nucleoporin NUP98 causes selective changes in nuclear pore complex stoichiometry and function. Proc Natl Acad Sci USA 98, 3191– 3196 (2001).

38. Hülsmann, B. B., Labokha, A. A. & Görlich, D. The Permeability of Reconstituted Nuclear Pores Provides Direct Evidence for the Selective Phase Model. Cell 150, 738– 751 (2012).

39. Frey, S. & rlich, D. G. O. FG/FxFG as well as GLFG repeats form a selective permeability barrier with self-healing properties. 28, 2554–2567 (2009).

40. Schmidt, H. B. & Görlich, D. Nup98 FG domains from diverse species spontaneously phase-separate into particles with nuclear pore-like permselectivity. eLife 4, 6281–30 (2015).

41. Deshmukh, L. et al. Structure and dynamics of full-length HIV-1 capsid protein in solution. J. Am. Chem. Soc. 135, 16133–16147 (2013).

42. Pornillos, O., Ganser-Pornillos, B. K., Banumathi, S., Hua, Y. & Yeager, M. Disulfide bond stabilization of the hexameric capsomer of human immunodeficiency virus. Journal of Molecular Biology 401, 985–995 (2010).

43. Faysal, K. M. R. et al. Pharmacologic hyperstabilisation of the HIV-1 capsid lattice induces capsid failure. bioRxiv 2022.09.21.508807 (2022). doi:10.1101/2022.09.21.508807

44. Song, Y. et al. Importin KPNA2 confers HIV-1 pre-integration complex nuclear import by interacting with the capsid protein. Antiviral Res 200, 105289 (2022).

45. Fernandez, J. et al. Transportin-1 binds to the HIV-1 capsid via a nuclear localization signal and triggers uncoating. Nature Microbiology 4, 1840–1850 (2019).

46. Krishnan, L. et al. The requirement for cellular transportin 3 (TNPO3 or TRN-SR2) during infection maps to human immunodeficiency virus type 1 capsid and not integrase. J. Virol. 84, 397–406 (2010).

47. Valle-Casuso, J. C. et al. TNPO3 is required for HIV-1 replication after nuclear import but prior to integration and binds the HIV-1 core. J. Virol. 86, 5931–5936 (2012).

48. Beck, M. et al. Nuclear pore complex structure and dynamics revealed by cryoelectron tomography. Science 306, 1387–1390 (2004).

49. Brass, A. L. et al. Identification of host proteins required for HIV infection through a functional genomic screen. Science 319, 921–926 (2008).

50. König, R. et al. Global analysis of host-pathogen interactions that regulate early-stage HIV-1 replication. Cell 135, 49–60 (2008).

51. Zhou, H. et al. Genome-scale RNAi screen for host factors required for HIV replication. Cell Host & Microbe 4, 495–504 (2008).

52. Yeung, M. L., Houzet, L., Yedavalli, V. S. R. K. & Jeang, K.-T. A genome-wide short hairpin RNA screening of jurkat T-cells for human proteins contributing to productive HIV-1 replication. J. Biol. Chem. 284, 19463–19473 (2009).

53. Di Nunzio, F. et al. Human Nucleoporins Promote HIV-1 Docking at the Nuclear Pore, Nuclear Import and Integration. PLoS ONE 7, e46037–15 (2012).

54. Di Nunzio, F. et al. Nup153 and Nup98 bind the HIV-1 core and contribute to the early steps of HIV-1 replication. Virology 440, 8–18 (2013).

55. Saito, H., Takeuchi, H., Masuda, T., Noda, T. & Yamaoka, S. N-terminally truncated POM121C inhibits HIV-1 replication. PLoS ONE 12, e0182434 (2017).

56. Guo, J. et al. The transmembrane nucleoporin Pom121 ensures efficient HIV-1 pre-integration complex nuclear import. Virology 521, 169–174 (2018).

57. Bayliss, R., Littlewood, T. & Stewart, M. Structural basis for the interaction between FxFG nucleoporin repeats and importin-beta in nuclear trafficking. Cell 102, 99–108 (2000).

58. Bayliss, R., Littlewood, T., Strawn, L. A., Wente, S. R. & Stewart, M. GLFG and FxFG nucleoporins bind to overlapping sites on importin-beta. J. Biol. Chem. 277, 50597– 50606 (2002).

59. Liu, S. M. & Stewart, M. Structural basis for the high-affinity binding of nucleoporin Nup1p to the Saccharomyces cerevisiae importin-beta homologue, Kap95p. Journal of Molecular Biology 349, 515–525 (2005).

60. Port, S. A. et al. Structural and Functional Characterization of CRM1-Nup214 Interactions Reveals Multiple FG-Binding Sites Involved in Nuclear Export. CellReports 13, 690–702 (2015).

61. Koyama, M., Hirano, H., Shirai, N. & Matsuura, Y. Crystal structure of the Xpo1p nuclear export complex bound to the SxFG/PxFG repeats of the nucleoporin Nup42p. Genes Cells 22, 861–875 (2017).

62. Gouveia, B. et al. Capillary forces generated by biomolecular condensates. Nature 609, 255–264 (2022).

63. Rasheedi, S. et al. The Cleavage and Polyadenylation Specificity Factor 6 (CPSF6) Subunit of the Capsid-recruited Pre-messenger RNA Cleavage Factor I (CFIm) Complex Mediates HIV-1 Integration into Genes. J. Biol. Chem. 291, 11809–11819 (2016).

64. Achuthan, V. et al. Capsid-CPSF6 Interaction Licenses Nuclear HIV-1 Trafficking to Sites of Viral DNA Integration. Cell Host & Microbe 24, 392–404.e8 (2018).

65. Sowd, G. A. et al. A critical role for alternative polyadenylation factor CPSF6 in targeting HIV-1 integration to transcriptionally active chromatin. Proc Natl Acad Sci U S A 113, E1054–63 (2016).

66. Francis, A. C. et al. HIV-1 replication complexes accumulate in nuclear speckles and integrate into speckle-associated genomic domains. Nature Communications 11, 3505–17 (2020).

67. Maertens, G. N. et al. Structural basis for nuclear import of splicing factors by human Transportin 3. Proc Natl Acad Sci U S A 111, 2728–2733 (2014).

68. Ambrose, Z. et al. Human immunodeficiency virus type 1 capsid mutation N74D alters cyclophilin A dependence and impairs macrophage infection. J. Virol. 86, 4708–4714 (2012).

69. Otsuka, S., Iwasaka, S., Yoneda, Y., Takeyasu, K. & Yoshimura, S. H. Individual binding pockets of importin-beta for FG-nucleoporins have different binding properties and different sensitivities to RanGTP. Proc Natl Acad Sci U S A 105, 16101–16106 (2008).

70. Blanco-Rodriguez, G. et al. Remodeling of the Core Leads HIV-1 Preintegration Complex into the Nucleus of Human Lymphocytes. J. Virol. 94, (2020).

71. Guedán, A. et al. HIV-1 requires capsid remodelling at the nuclear pore for nuclear entry and integration. PLoS Pathog 17, e1009484 (2021).

72. Márquez, C. L. et al. Kinetics of HIV-1 capsid uncoating revealed by single-molecule analysis. eLife 7, (2018).

## Methods References

1. Gagoski, D. et al. Gateway-compatible vectors for high-throughput protein expression in pro- and eukaryotic cell-free systems. J Biotechnol 195, 1–7 (2015).

2. Lau, D. et al. Fluorescence Biosensor for Real-Time Interaction Dynamics of Host Proteins with HIV-1 Capsid Tubes. ACS Appl Mater Interfaces 11, 34586–34594 (2019).

3. Lau, D. et al. Rapid HIV-1 Capsid Interaction Screening Using Fluorescence Fluctuation Spectroscopy. Anal. Chem. 93, 3786–3793 (2021).

4. Pornillos, O., Ganser-Pornillos, B. K., Banumathi, S., Hua, Y. & Yeager, M. Disulfide bond stabilization of the hexameric capsomer of human immunodeficiency virus. Journal of Molecular Biology 401, 985–995 (2010).

5. Piovesan, D., Monzon, A. M. & Tosatto, S. C. E. Intrinsic protein disorder and conditional folding in AlphaFoldDB. Protein Sci. 31, e4466 (2022).

6. Piovesan, D. et al. MobiDB: 10 years of intrinsically disordered proteins. Nucleic Acids Research 51, D438–D444 (2023).

7. Ng, S. C., Güttler, T. & Görlich, D. Recapitulation of selective nuclear import and export with a perfectly repeated 12mer GLFG peptide. Nature Communications 12, 4047–17 (2021).

8. Hagen, W. J. H., Wan, W. & Briggs, J. A. G. Implementation of a cryo-electron tomography tilt-scheme optimized for high resolution subtomogram averaging. Journal of Structural Biology 197, 191–198 (2017).

9. Kremer, J. R., Mastronarde, D. N. & McIntosh, J. R. Computer visualization of three-dimensional image data using IMOD. Journal of Structural Biology 116, 71–76 (1996).

